# Ancestral *versus* modern substrate scope in family-1 glycosidases

**DOI:** 10.1101/2024.04.11.589065

**Authors:** Luis I. Gutierrez-Rus, Dušan Petrović, Pascal Schneider, Katja Zorn, Gloria Gamiz-Arco, Rocío Romero-Zaliz, Ignacio Suarez-Martin, Leonardo De Maria, Francesco Falcioni, Valeria A. Risso, Martin A. Hayes, Jose M. Sanchez-Ruiz

**Author notes:** Correspondence aMartin A. Hayes. Discovery Sciences, BioPharmaceuticals R&D AstraZeneca, 431 50 Gothenburg, Sweden. Jose M. Sanchez-Ruiz. Departamento de Quimica Fisica. Facultad de Ciencias, Unidad de Excelencia de Quimica Aplicada a Biomedicina y Medioambiente (UEQ), Universidad de Granada, 18071 Granada, Spain. **Correspondence for manuscript submission:** Jose M. Sanchez-Ruiz. Departamento de Quimica Fisica. Facultad de Ciencias, Campus Fuentenueva s/n. Universidad de Granada, 18071 Granada, Spain.

## Abstract

Experimental studies support that protein engineering based on ancestral sequence reconstruction often leads to variants with biotechnologically useful biomolecular properties. These may include high stability, enhanced conformational flexibility and a modified catalysis range. Carbohydrate-active enzymes have numerous applications related with the degradation and synthesis of carbohydrates and glycoconjugates. Here, we explore how ancestral reconstruction may impact substrate scope in glycosidases, highly diverse enzymes that catalyze the hydrolysis of glycosidic bonds in all living cells and find applications as catalysts of the reverse reaction. To this end, we screen a library of ∼500 potential glycosidase substrates for degradation by both, a modern family-1 glycosidase from *Halothermothrix orenii* and a putative ancestral family-1 glycosidase derived from sequence reconstruction at a bacterial-eukaryotic common ancestor. The modern enzyme is the better catalyst for most substrates. But the ancestral glycosidase is more efficient with flavonoid glycosides bearing large aglycon moieties. Analysis of the catalytic parameters for a selected set of substrates, alongside analysis of the library data using a supervised learning algorithm, support the hypothesis that the modern enzyme tends to become less catalytically efficient with increasing substrate size, while this trend is not observed for the ancestral glycosidase. Molecular simulations support that the ancestral catalysis pattern is linked to the existence of a highly flexible region of the ancestral structure and a cavity capable of accommodating large aglycons. Our results provide guidelines for the engineering of enzymes for the synthesis and hydrolysis of large glycoconjugates.

## 1 | INTRODUCTION

Some 60 years ago, Pauling and Zuckerkandl proposed that plausible approximations to the sequences of proteins belonging to extinct organisms can be derived from comparative analyses of the known sequences of their modern descendants (Pauling & Zuckerkandl, 1963). Many proteins encoded by reconstructed ancestral sequences have been prepared and characterized in the last ∼25 years (Risso et al., 2013; Benner et al., 2017; Harms & Thornton, 2010; Merkl & Sterner, 2016; Devamani et al., 2016; Gumulya & Gillam, 2017; Risso et al., 2017; Gumulya et al., 2018; Spence et al., 2021; Selberg et al., 2021; Thomson et al., 2022; Nicoll et al., 2023; Jones et al., 2024). “Resurrected” ancestral proteins have been used to address important issues in molecular evolution. Furthermore, in a substantial number of cases, they have been found to display biomolecular properties that are useful in biotechnological applications. These include not only high stability, which plausibly reflects a hot start for life, but also enhanced conformational diversity and a modified catalysis range.

Conformational diversity is known to contribute to evolvability, *i.e.,* to the capability of generating new functionalities (James & Tawfik, 2003; Tokuriki & Tawfik, 2009; Petrović et al., 2018) and may promote catalytic promiscuity, a desirable feature in enzymes intended for biotechnological application (Zeymer & Hilvert, 2018; Trudeau & Tawfik, 2019).

Living organisms synthesize a wide diversity of carbohydrates and glycoconjugates with a variety of essential functions. Correspondingly, they also synthesize a huge diversity of carbohydrate-active enzymes, mostly glycoside hydrolases (glycosidases) and glycosyltransferases. Glycosidases are highly efficient enzymes that accelerate the rate of hydrolysis of glycosidic bonds by up to 17 orders of magnitude (Zechel & Winthers, 2000).However, they can also catalyze the reverse reaction under conditions that thermodynamically favor synthesis, such as high substrate concentrations of substrates, the utilization of activated substrates or use of cosolvents to reduce water concentration (Grunwald, 2018). Glycosidases can thus be used to couple sugars with glycosyl acceptors (*i.e.,* aglycons) yielding glycoconjugates with a diversity of potential applications (Grunwald, 2018).

Herein, we explore how ancestral reconstruction can impact substrate scope in glycosidases. This is a challenging goal since modern glycosidases are known to be promiscuous regarding the glycosyl acceptor (the aglycon), even if each glycosidase is typically rather specific for the glycosyl donor (the sugar) and the linkage generated or cleaved (α or β). Our approach thus relies on the screening of a large library of ∼500 potential glycosidase substrates for degradation by a modern glycosidase and a resurrected ancestral glycosidase. The modern protein we selected is a family-1 glycosidase from the halothermophilic organism *Halothermothrix orenii*. The ancestral protein selected is a putative ancestor of both eukaryotic and bacterial family-1 glycosidases that we have previously characterized in detail (Gamiz-Arco et al., 2021). Both enzymes have similar stabilities, as measured by the denaturation temperature values, and share the TIM-barrel fold. Yet, the ancestral protein display considerably enhanced conformational flexibility in some regions of the 3D-structure, as shown by molecular dynamics simulations, proteolysis experiments, and missing regions in electronic density maps derived from X-ray crystallography (Gamiz-Arco et al., 2021).

Library screening and subsequent analyses revealed a trend for the modern enzyme to become less catalytically efficient with increasing substrate size. This trend, however, is not observed in the ancestral protein. Actually, compared with the modern protein, the ancestral glycosidase is a much more efficient catalyst for the hydrolysis of large flavonoid glycosides (Figure 1). Molecular simulations provide a simple explanation for this result, revealing a cavity capable of accommodating large aglycon moieties in a conformationally flexible region of the ancestral protein structure. Overall, our results and analyses provide guidelines for the engineering of enzymes for the synthesis and hydrolysis of large glycoconjugates.

**FIGURE 1.**
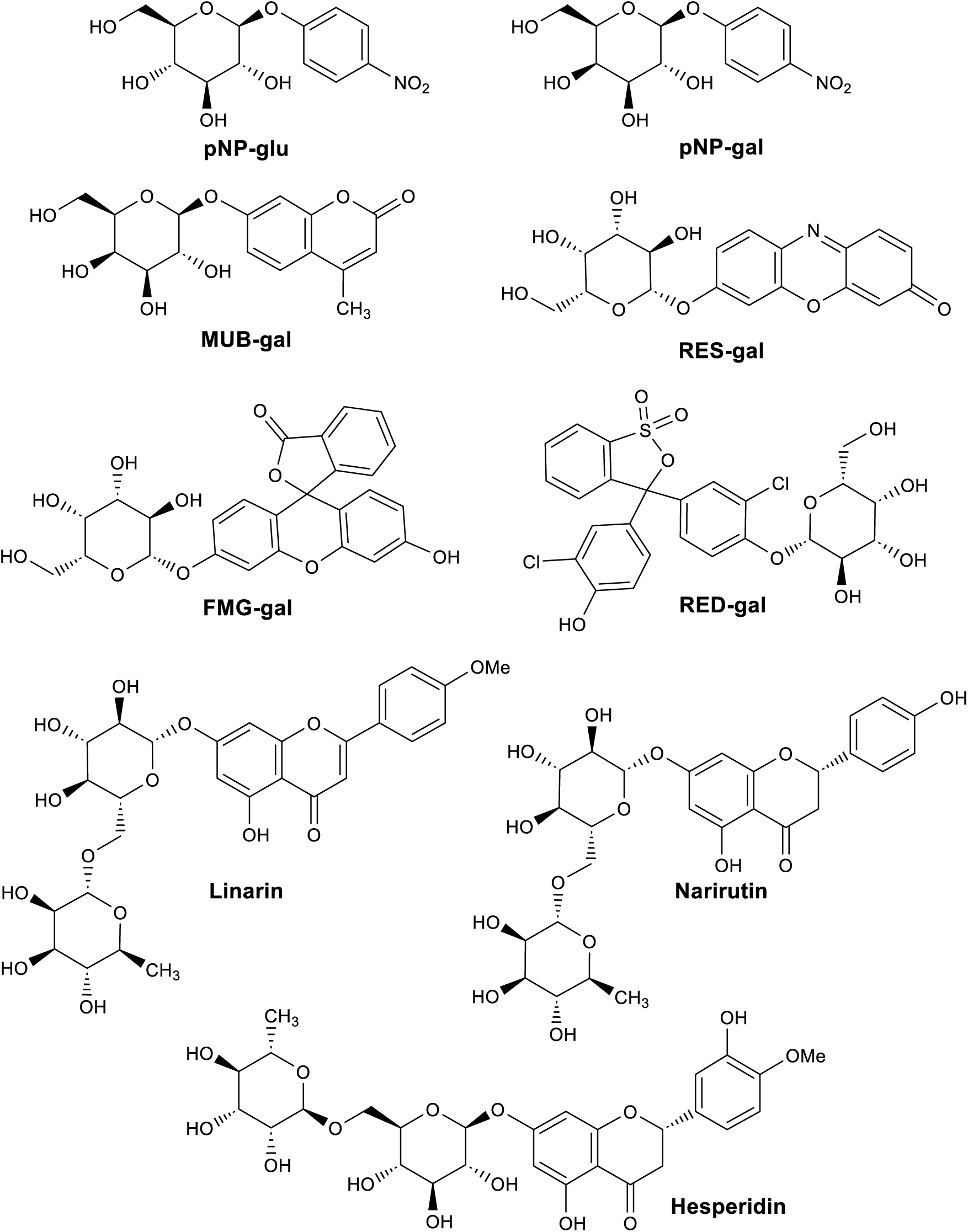
Glycosidase substrates discussed in detail in this work. pNP-glu: 4-nitrophenyl-b-D-glucopyranoside. pNP-gal: 4-nitrophenyl-b-D-galactopyranoside. MUB-gal: 4-Methylumbelliferyl β-D-galactopyranoside. RES-gal: Resorufin β-D-glucopyranoside. FMG-gal: Fluorescein mono-β-D-galactopyranoside. RED-gal: Chlorophenol Red-β-D-galactopyranoside. Glycoside flavonoids: linarin, hesperidin and narirutin.

## 2 | RESULTS

### 2.1 | Library screening for glycoside bond cleavage with the modern and ancestral proteins

We report an extensive comparison of the catalytic activity of an ancestral family-1 glycosidase with that of the modern family-1 glycosidase from *Halothermothrix orenii*. These two enzymes, ancestral and modern, display similar thermophilic stability and share a common TIM-barrel structure (Figure 2), but they display comparatively low sequence identity (58%) and noticeably different patterns of conformational flexibility (Figure 2), with the ancestral protein being clearly more flexible (Gamiz-Arco et al. 2021). Assessing glycosidase promiscuity is challenging, because these enzymes are widely diverse and are known to catalyze many different reactions. The current version of the CAZy database (The CaAZypedia consortium, 2018) lists 34 activities in family 1 alone and, additionally, the ancestral glycosidase may potentially catalyze some of the many reactions found in other families.

**FIGURE 2.**
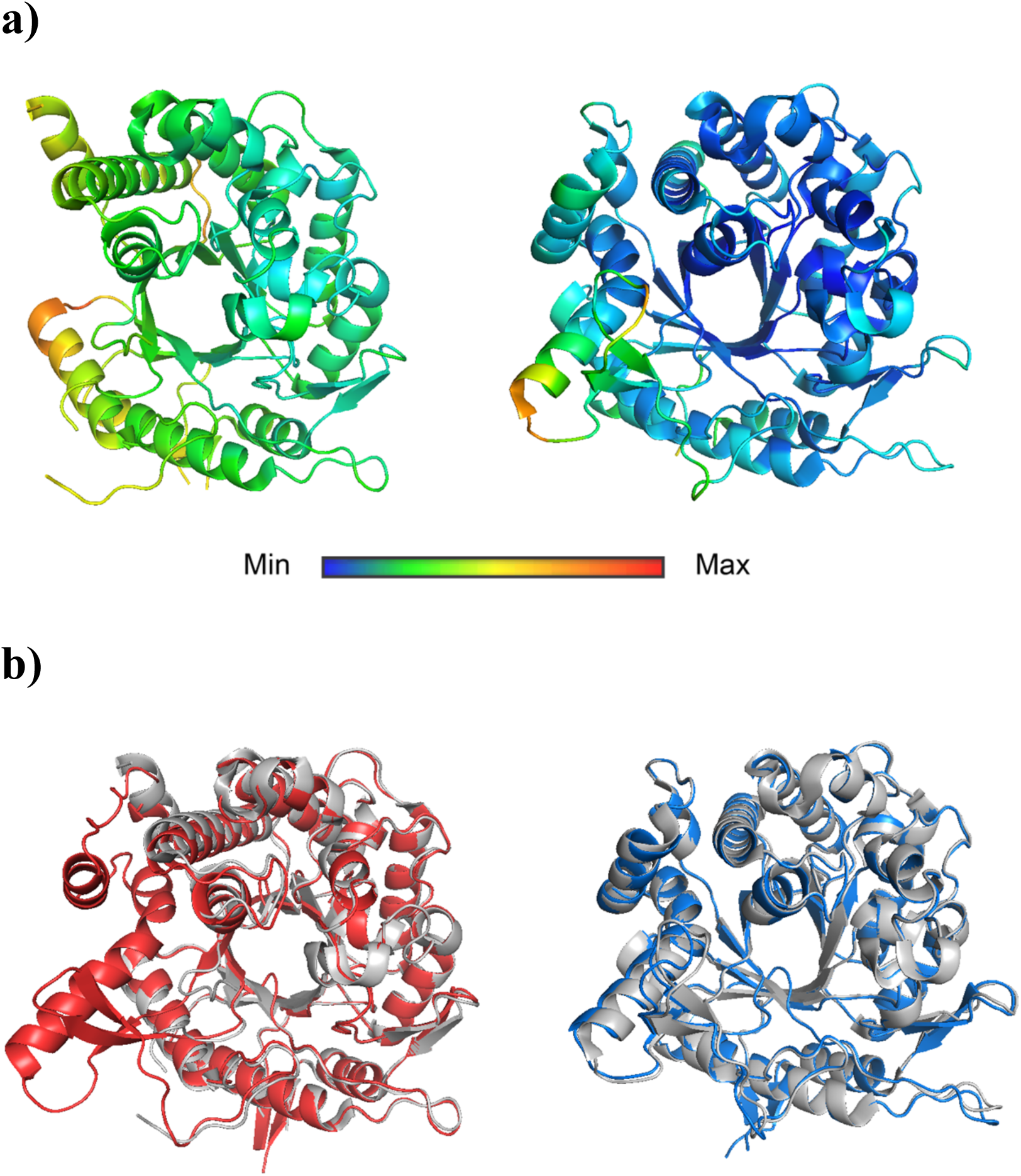
3D-structures of a putative ancestor of eukaryotic and bacterial family-1 glycosidases (the ancestral glycosidase) and the modern family-1 glycosidase from *Halothermothrix orenii* (the modern glycosidase). (a) Experimental structures derived from X-ray crystallography (PDB ID: 6Z1H for the ancestral glycosidase and PDB ID: 4PTV for the modern glycosidase). For illustration, structures have been colored according to normalized crystallographic B-factor values. The enhanced conformational flexibility of the ancestral proteins is actually supported by proteolysis experiments and by molecular dynamics simulations (Gamiz-Arco et al., 2021). Enhanced conformational flexibility in the ancestral protein is also revealed by missing regions in electronic density maps and, consequently, in the 3D-structure, as it is visually apparent. (b) Structures generated using Boltz-1 for the modern and ancestral glycosidase. For illustration, the predicted structures (blue for the modern enzyme and red for the ancestral enzyme) are superimposed with the experimental ones (grey). The agreement is excellent and the only noticeable difference is that Boltz-1 predicts the region of the ancestral structure that is missing from the experimental electronic density maps due to enhanced conformational flexibility.

Furthermore, glycosidases are already promiscuous to some extent regarding the aglycon moiety of the substrate (Grunwald, 2018). We thus performed a chemo-informatics screening of an internal chemical compound library at AstraZeneca, built using the following criteria: i) all the structures should contain a six-membered ring heterocycle (hexose and others), or potentially deoxy or otherwise modified sugar, ii) the glycosidic atom of the anomeric C1 of sugar was oxygen (and rarely nitrogen or sulphur), iii) the aglycon could either be another sugar or another organic molecule. As a result, ∼500 compounds were selected for experimental screening.

The main goal of the experimental screening was to identify “difficult” reactions with low levels of catalysis exhibited by the modern enzyme and higher levels of catalysis by the ancestral one. Therefore, comparatively high protein concentration (∼20 mM) and long incubation times (24 hours) were used for the in vitro substrate screening. The extent of reaction was assessed by UPLC-mass spectrometry (see Experimental Section in Supporting Information for details). The results were as follows. One hundred and eighty-nine of the targeted substrates were not detected and no further work was carried out to optimize their detection. For 176 of the substrates no significant conversion into products was observed, either with the modern enzyme or the ancestor. Fifty-five of the substrates appeared highly susceptible to glycosidase-catalyzed degradation and their complete conversion into products was observed with both, the modern and ancestral enzymes. For the remaining 60 substrates screened, significant conversion into products was observed with at least one enzyme but full conversion was not observed with both enzymes simultaneously.

For three substrates (narirutin, linarin and hesperidin) complete reaction after 24 hours was observed with the ancestral glycosidase, but not with the modern glycosidase. These three substates are glycosylated flavonoids with similar molecular structures (Figure 1). Interestingly, of the 34 activities listed for modern family-1 glycosidases in the CAZy database (The CaAZypedia consortium, 2018), only three appear to correspond unequivocally to the hydrolysis of flavonoid glycosides, and specifically the three activities involve substrates with anthocyanin or isoflavone as aglycons.

### 2.2 | Analysis of Michaelis-Menten catalytic parameters for a selected set of substrates (Figure 1)

We then carried out a more detailed characterization of the degradation of narirutin, linarin and hesperidin catalyzed by the modern and ancestral glycosidases. Kinetic experiments over a time scale of several hours followed by UPLC-mass spectrometry analysis confirmed that, in these three cases, hydrolysis leads to the non-glycosylated flavonoid and that the ancestral glycosidase is a more efficient catalyst than the modern glycosidase (Figure 3). Complete conversion of substrate to products is observed within hours with the ancestral glycosidase, but not with the modern glycosidase. Also, substantial hydrolysis was achieved in minutes using the ancestral glycosidase as catalyst but not with the modern glycosidase (Figure 3). Mass spectrometry was used to determine initial rates of reaction versus substrate concentrations (Figure 4a) and to determine kinetic parameters for the reactions (Figure 4b).

**FIGURE 3.**
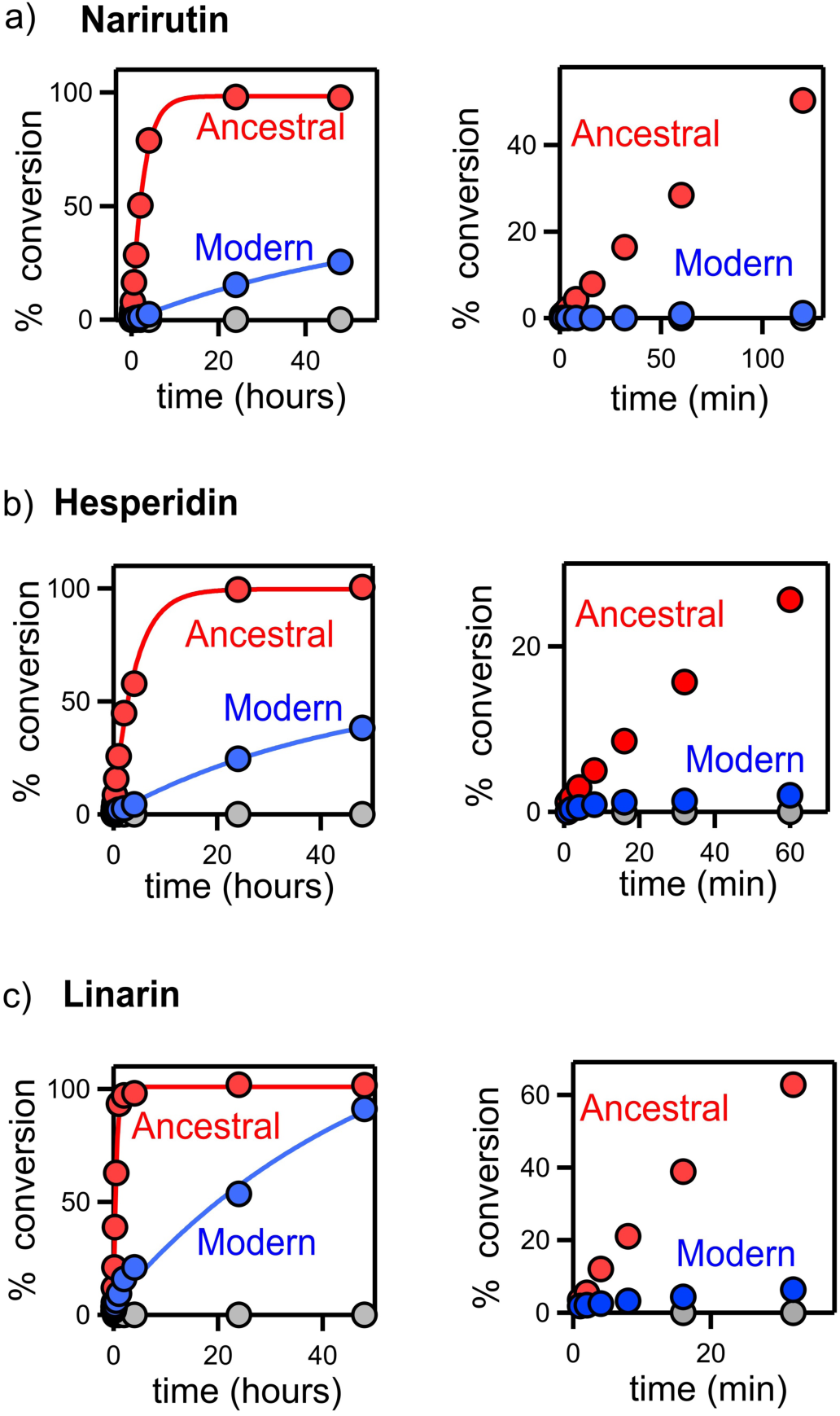
Kinetic of hydrolysis of flavonoid glycosides catalyzed by the ancestral glycosidase (red) and the modern glycosidase (blue). Blank experiments in the absence of enzyme are also shown (grey). Hydrolysis kinetics were followed based on the concentration of the flavonoid product, as determined by mass spectrometry. Note that the rate of the uncatalyzed reaction is negligible, in agreement with the known fact that glycosidic bonds display an extremely low rate of spontaneous hydrolysis (Zechel and Withers, 2000). The panels on the left show that, in all cases, full conversion into products is achieved in a time scale of hours with the ancestral enzyme as catalyst, but not with the modern enzyme. The panels on the right are blow-ups of the low time region and show that that significant hydrolysis is achieved in a time scale of minutes with the ancestral glycosidase as catalyst but not with the modern glycosidase.

**FIGURE 4.**
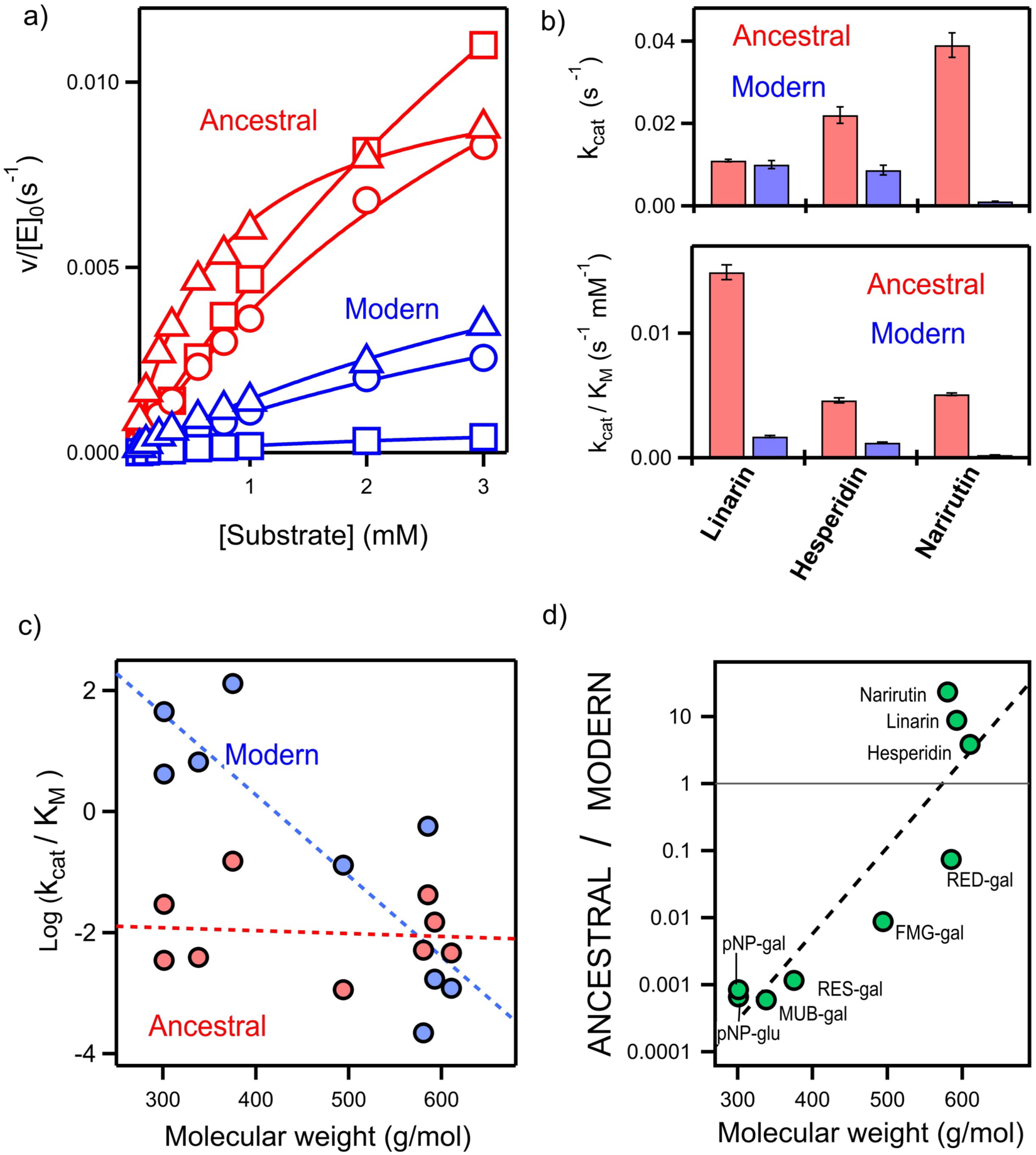
Catalytic parameters for glycoside hydrolysis. (a) Profiles of rate of hydrolysis *versus* substrate concentration for the hydrolysis of the flavonoid glycosides linarin (triangles), hesperidin (circles) and narirutin (squares), as catalyzed by the ancestral glycosidase (red) and the modern glycosidase (blue). [E]_0_ and v stand for the total enzyme concentration and the initial reaction rate. Continuous lines represent the best fits of the Michaelis-Menten equation to the experimental data. (b) Values of the catalytic rate constant (k_cat_) and catalytic efficiency (k_ca_t/K_M_) for the hydrolysis of flavonoid glycosides derived from the fittings shown in (a). Error bars represent standard deviations from the fittings. (c) Plot of logarithm of catalytic efficiency *versus* substrate molecular weight for the hydrolysis of all the substances shown in Figure 1, as catalyzed by the ancestral glycosidase (red) and the modern glycosidase (blue). The dashed lines are meant to guide the eye to the general trends. (d) Plot of the ancestral over modern catalytic efficiency ratio for all the substrates shown in Figure 1. The dashed line is meant to guide the eye to the general trend.

The values for the ancestral and modern catalytic efficiencies for the hydrolysis of the flavonoid glycosides are compared in Figure 4c with the values previously determined (Gamiz-Arco et al. 2021) for the hydrolysis of pNP-glu, pNP-gal, MUB-gal, RES-gal, FMG-gal and RED-gal (Figure 1). The comparison reveals patterns consistent with those expected for a substrate generalist enzyme (in the case of the ancestral glycosidase) and for a specialized enzyme (in the case of the modern glycosidase). The values for the modern glycosidase span about 6 orders of magnitude and show a clear trend towards decreased catalytic efficiency with increasing substrate molecular weight while the pattern observed for the ancestral glycosidase is completely different (Figure 4c). Catalytic efficiency values are similar with the ancestral enzyme (for up to two orders of magnitude) for the various substrates in Figure 1 and no clear trend with molecular weight is apparent. As a result, while the ancestral glycosidase is a much less efficient catalyst than the modern enzyme with typical family-1 glycosidase substrates, e.g. pNP-glu, pNP-gal, the two enzymes display similar levels of catalysis with the larger substrates in Figure 1, as shown in Figure 4c. The ancestral glycosidase is the more efficient catalyst for flavonoid glycoside substrates (Figure 4d).

### 2.3 | Molecular simulations on the interaction of selected glycosides (Figure 1) with the modern and ancestral glycosidases

To further explore the factors that determine the different patterns of catalysis for the ancestral and the modern glycosidases, we have carried out computer simulations on substrate-protein interactions using the Boltz-1, a deep-learning open-source program that predicts 3D-structures for biomolecular complexes with alphafold3 accuracy. The enzyme sequences given as input to the program are listed in the supplementary material as well as the SMILES strings for the substrates (Table S1). In the absence of substrates bound, the structures predicted by Boltz-1 are in excellent agreement with the experimental X-ray structures of both the modern glycosidase and the ancestral glycosidase (Figure 2). It must be noted, however, that a comparatively large region is missing from the experimental ancestral structure, while it is obviously present in the prediction. This reflects missing experimental electronic density linked to much enhanced conformational flexibility in the aforementioned region, as we have previously discussed in detail.

Representative examples of the 3D-structures for the glycosidase-substrate complexes predicted by Boltz-1 are shown in Figure 5a (additional illustrative examples are given in Figure S1). Several common features are apparent. First, the glycosidic bond to be cleaved in the substrate molecule is reasonably close to the catalytic carboxylic acid residues of the glycosidases’ active sites. Second, the aglycon moiety of the docked substrate is accommodated in a cavity formed in which there is an abundance of hydrophobic residues with mid to short side chain lengths, while in the modern scaffold this cavity is not present, as the space is mainly occupied by aromatic residues. This explains the differences between ancestral and modern scaffold” This is obviously more clearly seen in the structures with the docked flavonoid glycosides (narirutin, linarin, hesperidin), which are large molecules. The relevant point here is, however, that the aglycon moiety binds in the region of the 3D-structure that is known from previous studies to be conformational flexile. This result supports enhanced conformational flexibility as one of the molecular determinants of the improved efficiency of the ancestral enzyme towards glycoside flavonoids, as compared with its modern counterpart.

**FIGURE 5.**
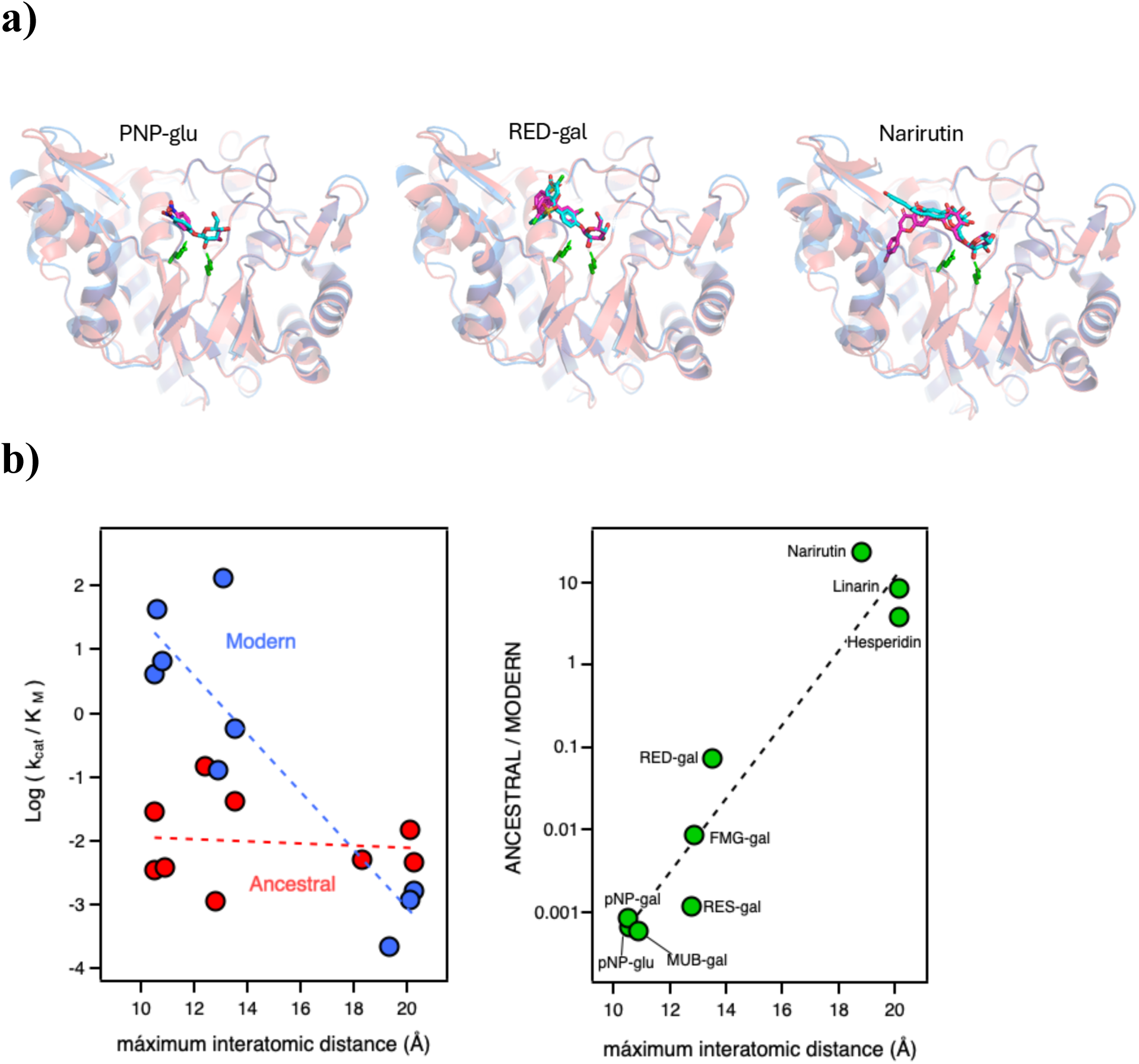
Molecular simulations on the interactions of glycoside substrates with the modern and ancestral glycosidases. (a) Protein structures with substrates docked, as predicted by Boltz-1. Modern is shown in blue and ancestral is shown in red Here we show some representative examples; additional illustrative predictions can be found in Figure S1. The catalytic carboxylic acid residues at the glycosidase active site are shown. (b) The panel at the left is a plot of logarithm of catalytic efficiency *versus* maximum inter-atomic distance for the docked substrates for the hydrolysis of all the substances shown in Figure 1, as catalyzed by the ancestral glycosidase (red) and the modern glycosidase (blue). The dashed lines are meant to guide the eye to the general trends. The panel at the right is a plot of the ancestral over modern catalytic efficiency ratio *versus* maximum inter-atomic distance for the docked substrates for all the substrates shown in Figure 1. The dashed line is meant to guide the eye to the general trend.

In addition, the Boltz-1 simulations shed light on the dependence of catalysis with substrate size. Clear trends with substrate molecular weight are thus observed in Figures 4c and 4d. However, it is apparent that the correlations are far from being perfect. The ancestral versus modern catalytic efficiency ratio for the flavonoid glycosides (narirutin, linarin, hesperidin) is thus about two orders of magnitude higher than that for the RED-gal substrate, even though the 4 compounds have similar molecular weights. However, in our Boltz-1 simulations RED-gal (and also FMG-gal) appears to occupy smaller regions of the protein, as compared with the glycoside flavonoids, despite having similar molecular weights (in particular for RED-gal). The diminished interaction of FMG-gal and RED-gal with the proteins is obviously due to a different shape of the aglycon moiety, which is determined by the presence of a tetrahedral spiro sp^3^ carbon connecting several rings. Clearly, the Boltz-1 simulations confirm that size is a main determinant of catalysis in these systems, *i.e.*, the but also support that molecular weight is not a perfect metric of size in this context. As an alternative metric, we have calculated the maximum inter-atomic distance for the docked substrates. Using this metric of size, instead of molecular wight significantly improves the correlations involving catalytic efficiency (see Figures 5b and 5c).

### 2.4 | Computational analysis of glycoside-binding cavities in the modern and ancestral glycosidases

To further explore the molecular determinants behind the different catalytic behaviours of the modern and the ancestral glycosidase pockets/cavities were identified by using Fpocket software tool (Le Guilloux et al., 2009) as part of the CaverWeb suite (Marques et al., 2025). Precited structures for both glycosidases were used as inputs in CaverWeb and pocket analysis was performed using default parameters. Residues that define the pockets were also identified in CaverWeb and subsequently confirmed by visual inspection. A cavity was apparent in the predicted structures of both, the modern and the ancestral glycosidases matching the glycosidase active site at which the sugar moiety of the glycoside is found to bind in the Boltz-1 simulations (Figure 6). However, CaverWeb finds a cavity matching the large aglycon moiety of the flavonoid glycosides only in the predicted ancestral structure (Figure 6). The absence of such a cavity, as probed by CaverWeb, in the modern structure is likely linked to the presence of a number of bulky side chains which are replaced by less bulky side-chains in the ancestral structure.

**FIGURE 6.**
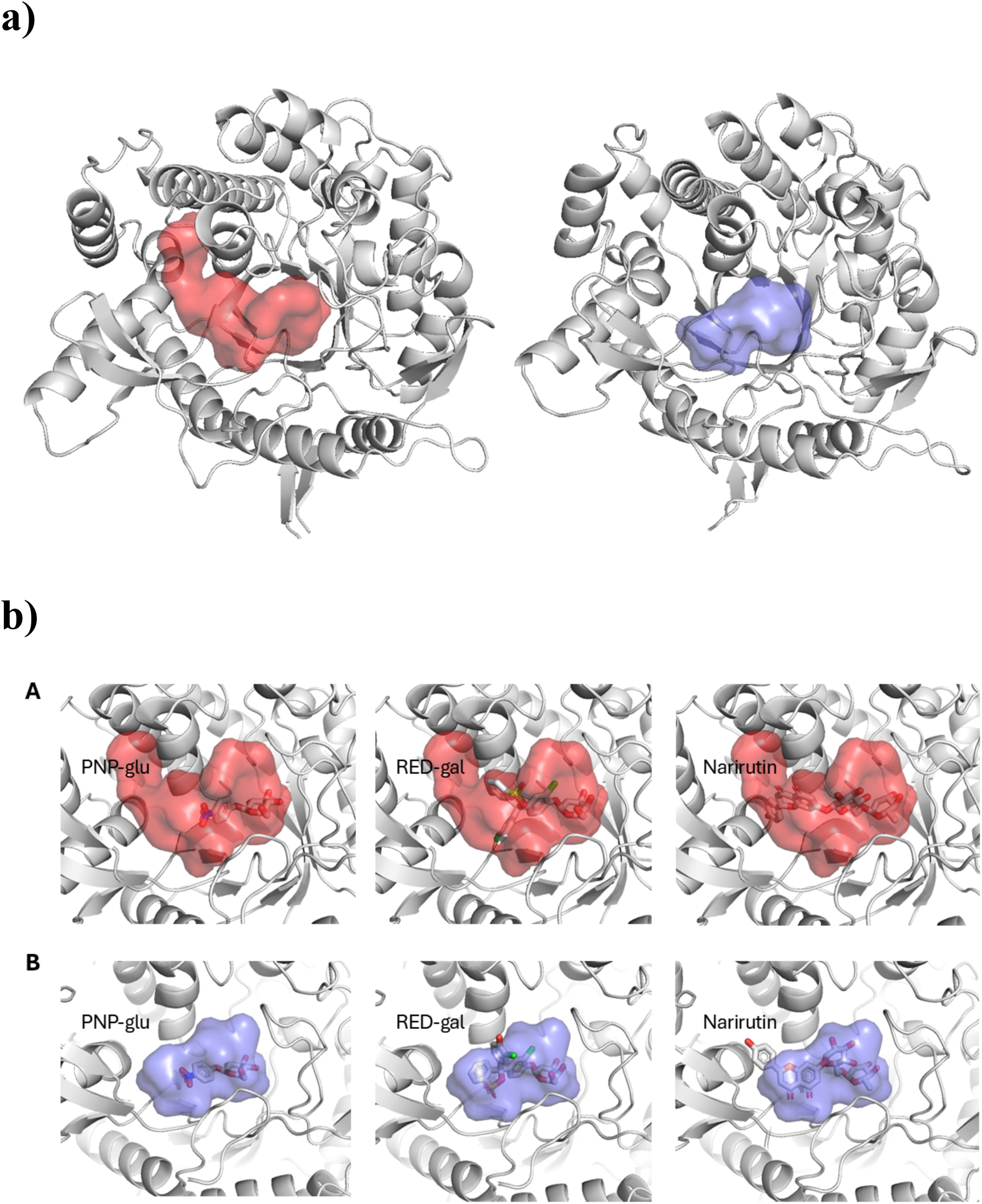
Analysis of cavities in the structures of the. The cavity in the modern protein is shown in blue and the cavity in the ancestral protein is shown in red. (b) Cavities and docked substrates for several illustrative examples. Cavity color code is the same as in panel a. Note that narirutin, a substrate with a large aglycon moiety, does not fit the cavity.

Boltz-1calculations on the binding free energies for the substrates of Figure 1 yield results generally consistent with the different cavity patterns in the modern and ancestral glycosidases structures. For the substrates of smaller size among those in Figure 1, the free substrates, energies of binding to the modern protein are similar to the free energies of binding to the ancestral proteins (Figure 7A). On the other hand, a trend towards lower binding energies with the ancestral protein is observed with the larger substrates (Figure 7A), thus supporting the enhanced ancestral capability to accommodate large substrates. It is to be noted that each value shown in Figure 7 is the average of the results obtained in computations performed with 10 different structures, as predicted by Boltz-1 with the scatter of the ten calculated values (standard deviation from the average) being shown by the associated bars in Figure 7. Interestingly, a trend towards larger scatters is observed for the larger substrates, in particular with the data corresponding to the binding to the modern glycosidase. Plausibly, binding to the modern structure, in which a cavity that can accommodate large aglycons is not apparent, is more susceptible to small structural changes in the neighbourhood of the binding site.

**FIGURE 7.**
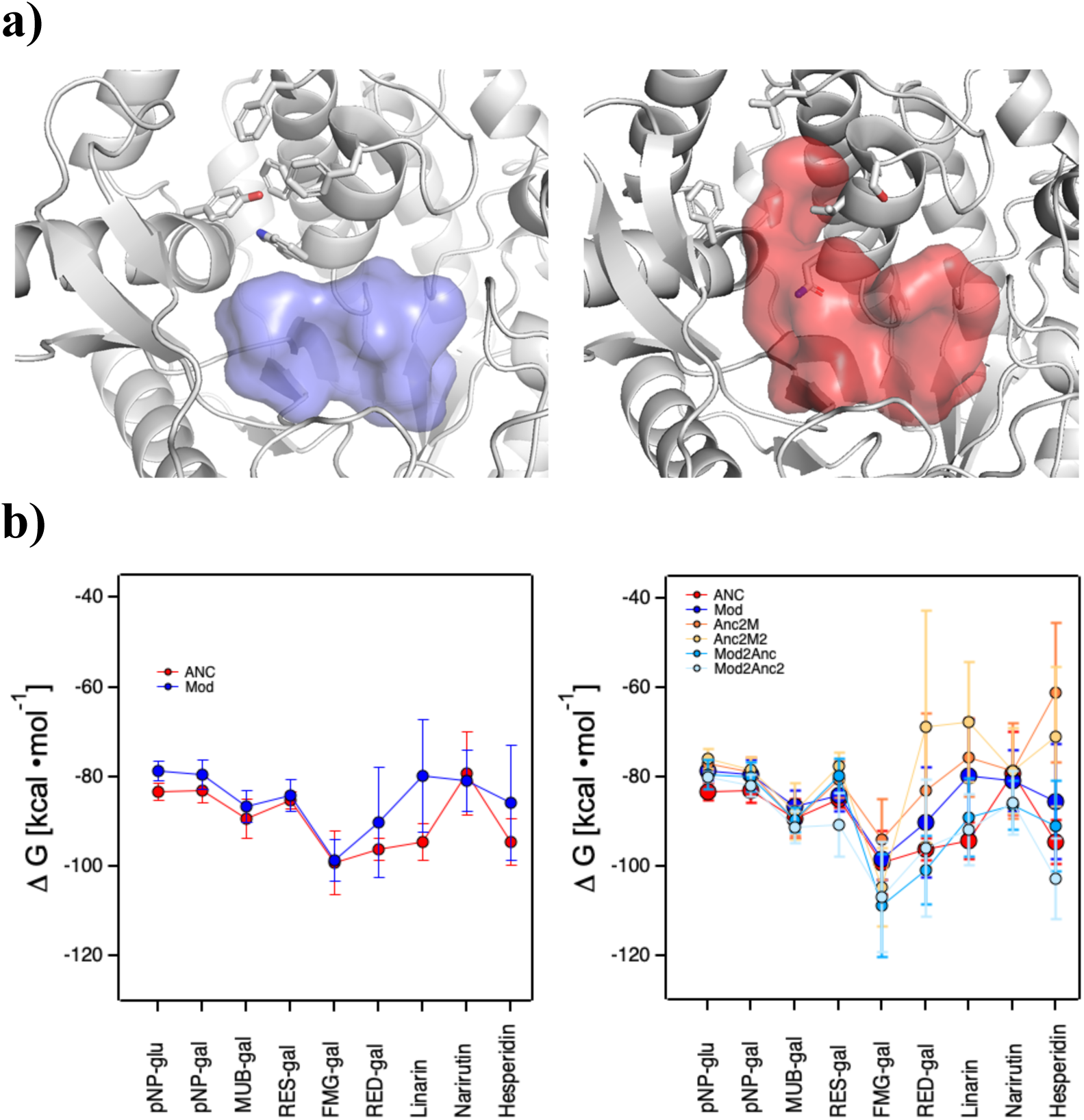
Computational analyses into the mutational effects on glycoside binding. a) 3D-structures as predicted by Boltz-1 showing residues that corresponding to the cavity in the ancestral protein structure that can accommodate large aglycin moieties (red) and the corresponding residues in the modern structure. b) Data for the binding of the several substrates to the modern and ancestral glycosidases. Boltz-1 was used to calculate free energy changes for the binding of the substrates in Figure 1 to the wild-type forms and several variants of the modern and ancestral glycosidases. Left: data for the binding of the several substrates to the modern protein and the ancestral protein. Right: data for the binding of the several substrates to the modern protein and to two multiple-mutation variants of the modern protein. The mutations introduced are as follows. Mod2Anc: W168N, F172V, E173L, A176L, F187L, T224S, I246F, S297T, M299N; Mod2Anc2: W168N, F172V, E173L, A176L, F177S, F187L, T224S, L242R, I246F, E262M, Y267G, S297T, M299N. Also in the right panel: Data for the binding of the several substrates to the ancestral protein and to two multiple-mutation variants of the ancestral protein. The mutations introduced are as follows. Anc2M: N173W, V177F, L178E, L181A, L192F, S229T, F251I, T305S, N307M. Anc2M2: N173W, V177F, L178E, L181A, S182F, L192F, S229T, R247L, F251I, M267E, G272Y, T305S, N307M. In all cases, the average of 10 calculations with 10 different structures predicted by Boltz-1 is given. The bars represent the scatter of the 10 calculations, as measured by the standard deviation from the average

### 2.5 | Computational analyses into mutational effects on glycoside binding

The computational analyses on binding cavities reported in the previous section suggest that increased efficiency of the ancestral glycosidase towards flavonoid glycosides as compared with its modern counterpart is linked to the presence of a suitable cavity capable to accommodate comparative large aglycons, which is likely promoted by enhanced conformational flexibility. The possibility arises then that suitable mutational changes could generate the cavity in the modern structure and, more generally, that back-to-the-ancestor engineering can be used to engineer binding of hydrolysis of large-aglycon glycosides. Here, we use structural modelling and computational prediction on binding affinities to explore this possibility.

Visual comparison of the ancestral and modern structures may be used to suggest combinations of back to the ancestor mutational changes that could make the modern protein more “ancestral” in terms of the presence of a cavity capable of accommodating large aglycons. Conversely, the corresponding ancestral-to-modern amino acid replacements could make the ancestral protein more “modern” in terms of the closing of the cavity. We have explored two such combinations of mutations, specifically those involving positions [173, 177, 181, 192, 229 251, 302] and positions [247, 267, 173, 177, 178, 181, 192, 229, 251, 302, 304], in both cases using the ancestral protein numbering). Boltz-1 calculations on the binding energies for all the substrates of Figure 1 show that the back-to-the ancestor mutations in the modern glycosidase promote binding of the larger substrates, as more negative binding affinities are obtained (see Figure 7B and the legend to the figure for the specific mutations introduced). Conversely, the corresponding ancestor-to-modern mutations lead to a less favourable binding affinities when introduced in the ancestral protein (see Figure 7C and the legend to the figure for the specific mutations introduced).

In order to further explore the possibility of cavity engineering for large aglycon binding, we carried out Boltz-1 calculations of binding affinities for single-mutant variants of the modern the ancestral glycosidases. Each variant of the modern protein included a single back-to-the-ancestor mutation, while each variant of the ancestral protein included a single ancestral-to-modern mutation. Variants with mutations at positions close to the cavity that binds the aglycon in the ancestral structure were studied. The results show that mutational effects on the binding free energies tend to be more prominent for the substrates with large aglycon moieties. However, a small set of back-to-the-ancestor replacements responsible for the more favourable binding calculated for the combinations of mutations of Figure 7. This is not surprising, given that a single mutation is unlikely to generate by itself an intramolecular environment capable to accommodate large aglycon moieties. In fact, generation of a capability to accommodate large aglycons will likely require the cooperative and possibly synergistic effects of several mutations. Such epistatic effects are difficult to predict and the reliability of computations involving large numbers of mutations is thus limited. Our computational results do support that cavity engineering for large aglycon accommodation is feasible, although it appears that the screening of a combinatorial library of the targeted mutations would probably provide more efficient way to achive this goal.

### 2.6 | Analysis of the library data using a supervised learning algorithm

The computational and kinetic analyses reported in the preceding sections suggest different trends relating to substrate size for the modern and the ancestral glycosidase. These different trends have been inferred, however, from the consideration of a limited set of 9 substrates (Figure 1) for which catalytic efficiency values are available. It is reasonable to ask to what extent these trends are general or, more specifically, whether the observed trend with substrate size is also apparent in the data for the library of several hundred compounds studied here. Addressing this question is not straightforward for two reasons. Firstly, we do not have catalytic efficiency values for the majority of the library compounds, and only a qualitative assessment of their susceptibility to degradation by the glycosidases in a 24 hour time period (Table S2).

That is, for each glycosidase, modern and ancestral, the compounds were assigned to the one of three classes a) “no degradation observed”, b) “partial degradation observed”, c) “full degradation observed”. Secondly, for only three of the compounds in the library, the flavonoid glycosides linarin, narirutin and hesperidin, the ancestral glycosidase was found to be more efficient that than the modern glycosidase, according to the qualitative classification. This makes it very difficult to use the screening to question the factors that make these glycosides better substrates for the ancestral protein compared to the modern one.

In view of the above, we decided to ask the opposite question. That is, we aimed at exploring which factors could make certain glycosides better substrates for the modern protein compared to the ancestral one. To this end, we first divided the library compounds in two discrete sets. Set 1 included all the compounds for which the modern protein was found by our assay to be more efficient than the ancestral protein; that is, set 1 includes all compounds for which the modern glycosidase achieves full degradation, while the ancestral glycosidase achieves only partial or no degradation, as well as compounds for which the modern glycosidase achieves partial degradation and no degradation is observed with the ancestral protein. Fifty compounds were included in Set 1. The remaining compounds, *i.e.,* the compounds for which the modern protein was not determined to be more efficient than the ancestral protein by our assay, were included in Set 2 (241 compounds). Next, molecular descriptors were selected to represent properties of the compounds that could conceivably impact their susceptibility to undergo glycosidase catalysed cleavage. These descriptors included molecular weight, of course, but also parameters related to molecular features of the aglycon and carbohydrate moieties. The original set of descriptors is provided in Table S3. Actually, a correlation analysis (Figure S2) was performed and a smaller set essentially uncorrelated descriptors was retained for subsequent analysis (Table S3). Finally, a decision tree (Quinlan, 1983; von Winterfeldt and Edwards, 1986) was defined using the reduced set of molecular descriptors, providing an interpretable classification framework which enable us to highlight the most relevant variables contributing to molecular classification in while keeping transparency in the decision-making process. The model was built using the Gini index (Gini, 1936) as the splitting criterion, which evaluates feature importance by measuring impurity reduction at each node. The approach was applied to the classification of the library compounds in the two sets described above, but it allowed the recursive partition of the descriptors space into clusters, facilitating the identification of patterns associated with the target classes. The decision tree obtained is shown in Figure 8a. It reveals a cluster with modern>ancestral preference at low substrate molecular weight (between 447 and 469 g/mol) together with clusters with the same preference for substrates with large numbers of heterocycles or carbocycles in the aglycon moiety.

**FIGURE 8.**
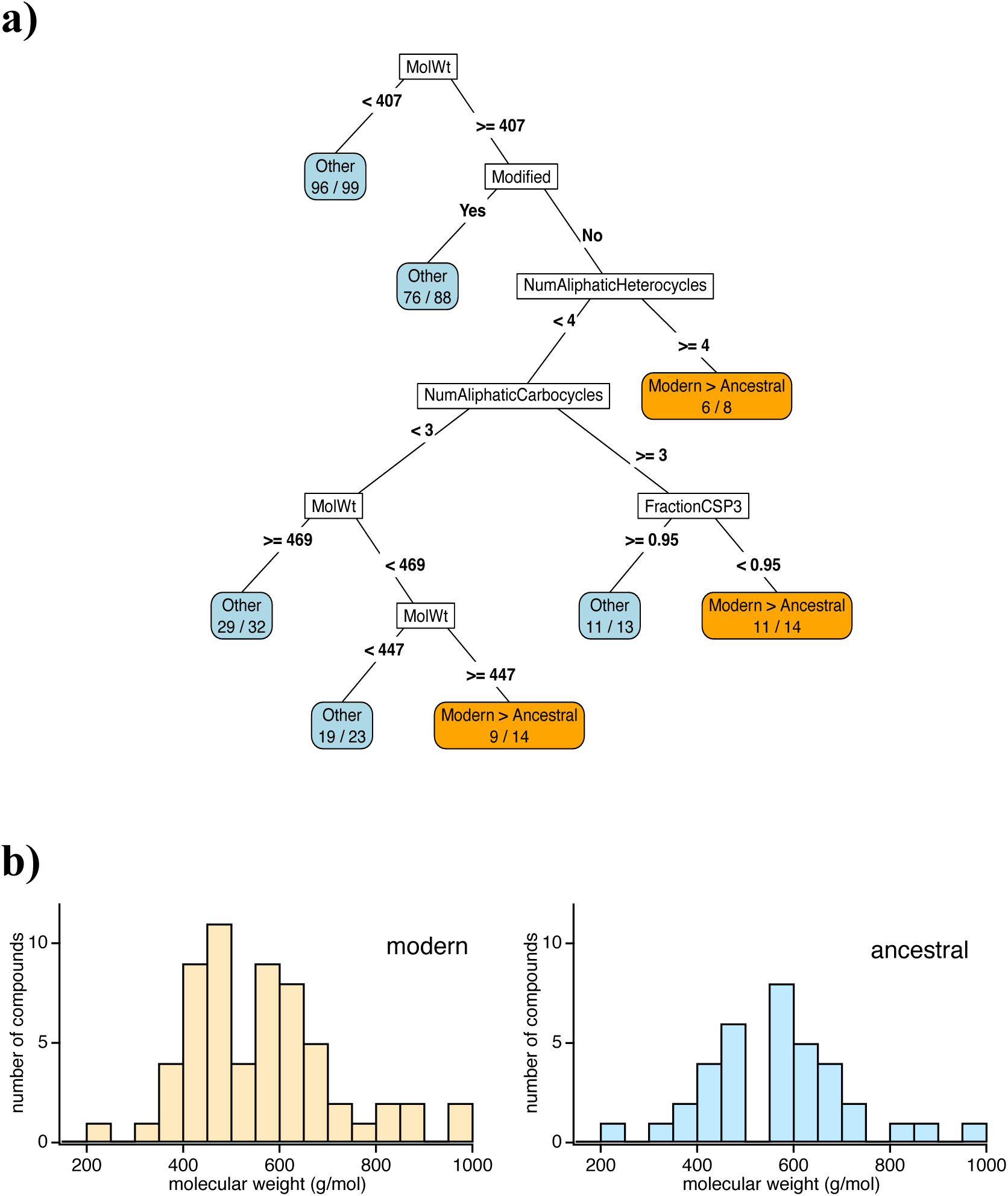
Analysis of library data using a supervised learning algorithm. Library compounds are divided in two disjoint sets, one including all the compounds for which the modern protein is found by our assay to be more efficient than the ancestral protein (Set 1: 50 compounds) and another set including all the compounds for which the modern protein is not found by our assay to be more efficient than the ancestral protein (Set 2: 241 compounds). A decision tree (**a**) was defined using a suitable set of molecular descriptors thus providing an interpretable classification framework. The terminology used for the descriptors is explained in Table S3. The analysis classifies the variants in several clusters, which are coloured according to the most represented Set. Clusters for which most of the compounds belong to Set 1 (modern enzyme is more efficient than the ancestral enzyme) are highlighted in orange. All other clusters are coloured in blue. For each cluster, the total number of compounds is shown at the right and the number of compounds belonging to the most represented set is shown at the left. (**b**) Molecular weight distributions for library compounds susceptible to hydrolysis by the modern enzyme (left) and the ancestral enzyme (right).

This result obtained from the decision-tree analysis are consistent with the molecular weight distributions for library compounds susceptible to hydrolysis by the modern enzyme and the ancestral enzyme (Figure 8b). The distribution peaks at about 450g/mol for the modern glycosidase and at about 600 g/mol for the ancestral glycosidase. Furthermore, the modern enzyme appears to be statistically more efficient than the ancestral enzyme for the substrates with the highest molecular weight within the screened library. One possibility is that the ancestral preference for larger glycosides is determined to a substantial extent by the size of the cavity present in the ancestral structure which can readily accommodate extended substrates up to around 600 g/mol (sections 2.4 and 2.5 above). Such preference, therefore, may not exist for much larger substrates. In this interpretation, there is a “goldilocks” substrate size for the ancestral enzyme corresponding to a molecular weight of about 600 g/mol. Yet, compounds with molecular weight much higher than 600 g/mol are underrepresented in the screened library and, consequently, discussions on the ancestral vs. modern trends at high molecular weight are necessarily speculative

## 3 | DISCUSION

The enhanced catalysis observed with the ancestral enzyme with flavonoid glycosides poses intriguing questions from an evolutionary context. Flavonoids are highly diverse secondary metabolites produced by fungi and land plants, often as glycosidic derivatives. Fungi diverged roughly 1.3 billion years ago, while land plants evolved from algae about 1 billion years ago (Kumar et al., 2022). On the other hand, our putative ancestral glycosidase derives from sequence reconstruction targeting a common ancestor of bacteria and eukaryotes, *i.e.*, a very ancient phylogenetic node (about 4 billion years). Under the, admittedly speculative, assumption that our putative ancestral glycosidase reproduces to some extent the properties of glycosidases that existed ∼4 billion years ago, it is somewhat puzzling that it hydrolyzes compounds that are commonly found in organisms that evolved much later. Of course, we cannot rule out that flavonoid glycosides had roles in early life and were already produced ∼4 billion years ago. Another possibility is simply that the ancestral glycosidases were generalists (Jensen, 1976; ÓBrien & Herschlag, 1999; Zou et al., 2015), capable to hydrolyze a diversity of glycoside substrates with various aglycons and that flavonoid glycosides simply resemble the actual ancestral substrates.

Beyond our more evolutionary interpretations, these results provide one experimental example in which ancestral reconstruction modulates glycosidase substrate scope towards glycosides bearing large aglycons. Furthermore, our analyses highlight the biomolecular features responsible for this modulation. Computer simulations on the interaction of flavonoid glycosides with both studied proteins indicate that the large aglycon moiety fits in a region of the 3D-structure which is conformationally flexible in the ancestral glycosidase, but not so in the modern glycosidase. Clearly, this biomolecular feature of the ancestral protein should facilitate substrate binding in catalytically competent conformations. Furthermore, structural analyses reveal a preformed cavity suitable for aglycon binding in the ancestral protein. The cavity appears to some extent be to some extent to be blocked by amino acid side chains in the modern protein, implying that a larger conformational change would be required in this case for substrate binding.

Glycosidases are typically specific for given glycosyl donors (the sugar) and the linkage generated (α or β) but are known to exhibit promiscuity towards the aglycon moiety of the substrate This is relevant since glycosidases can be used to catalyze the coupling of sugars with aglycons to yield glycoconjugates, which have a diversity of applications. Furthermore, in several cases, the glycoconjugates of interest may have large aglycons. For instance, cardiac glycosides are derived from steroid aglycones, and conjugating sugars to proteins has been used to increase half-life and in vaccine development (Grunwald, 2018). This work provides guidelines for the engineering of enzymes for the synthesis and degradation of large glycoconjugates. Our analyses support that rationally designed mutational replacements may contribute to generating a suitable cavity and that ancestral reconstruction engineering may lead to enhanced conformational diversity facilitating aglycon placement in such cavity through enhanced conformational sampling. Glycosidases are widely distributed and are currently classified into about 200 evolutionary distinct families (The CAZpedia Consortium, 2018). Although our results address family-1 glycosidases, it seems plausible that the general guidelines offered here could be applied to other glycosidase families or even to other carbohydrate-active enzymes, such as glycosyltransferases.

## 4. | MATERIALS AND METHODS

### 4.1 | Protein expression and purification

The ancestral and modern proteins studied in this work were prepared as previously described (Gamiz-Arco et al., 2021). Genes for each protein were cloned into a pET24b(+) vector with kanamycin resistance (GenScript) containing a 6xHis-tag located in the C-terminus of the protein sequence. Plasmids were transformed into *E. coli* BL21(DE3) cells following a standard heat shock transformation protocol. Protein expression was carried out in cultures with 1 L of LB media supplemented with kanamycin. Cells were grown at 37°C to reach an optical density of OD_600_=0.6 and then IPTG 0.4 mM was added to the cultures to induce protein overexpression during 16 hours of incubation at 37°C with agitation at 250 rpm. Cells were collected by centrifugation, resuspended in binding buffer (HEPES 20 mM, NaCl 500 mM, imidazole 20 mM) and sonicated while kept in ice. Sonication was performed in presence of protease inhibitors (cOmplete® EDTA-free Protease Inhibitor Cocktail, Roche) since the enhanced conformational flexibility of the ancestral protein makes it more susceptible to proteolysis. Proteins were purified by immobilized metal affinity chromatography using Ni-NTA columns (His GraviTrap Columns, Cytiva). Buffer exchange for each protein was performed by passing through PD-10 desalting columns prepacked with Sephadex G-25 resin (Cytiva) and pre-equilibrated with HEPES 50 mM pH 7.00 NaCl 150 mM. Finally, proteins were further purified by size exclusion chromatography using the same buffer and a HiLoad Superdex 200 pg preparative column. Central factions of the eluted peak were collected to ensure maximum purity and were concentrated for further experiments. Protein purity was checked by SDS-PAGE.

### 4.2 | Generation and preparation of libraries

Libraries of chemical compounds used as substrates to interrogate the enzymatic activity of both ancestral and modern glycosidases were designed and prepared as followed. First, a computational screening was performed using an internal chemo-informatics compound library from AstraZeneca according to a three-step criterium: I) A first screening was applied to select all compounds with molecular structures containing a hexose or pentose sugar, or potentially deoxy or otherwise modified sugar, II) the glycosidic atom at the anomeric C1 of the sugar was oxygen (or rarely via other atoms e.g. like nitrogen or sulphur), III) the aglycon could either be another sugar or another organic molecule. As a result, 480 different compounds were selected for high throughput screening enzymatic assays, for which 1 µl of each selected compound at concentrations of ∼1 mM dissolved in DMSO was dispensed into 384-well high recovery microplates (384 Well Microplate, PP, V-Bottom, Greiner Bio-One) using an Echo 555 Acoustic Liquid Handler (Beckman Coulter).

### 4.3 | Miniaturised high-throughput screening with Ultrahigh-Performance Liquid Chromatography coupled to Mass Spectrometry (UPLC-MS)

Reactions were performed on 12 µl scale in 384-well high recovery microplates covered with a silicon seal. Substrates from the compound library were previously diluted in HEPES 50 mM pH 7.00 NaCl 150 mM to a concentration of 0.7 mM by adding a volume of buffer into each microwell using an OT-2 Liquid Handling Robot (Opentrons). Substrate solutions were mixed by shaking the microplates at 1000 rpm for 5 minutes to ensure a correct homogenization. Then, 10 µl of the enzyme solution at concentrations of ∼1 mg/ml or ∼20 µM were dispensed into a new reaction microplate. For control reactions, HEPES buffer was added to the reaction plates instead of protein solution. Reactions were started by addition of 2 µl substrate solution from the library microplates to the reaction microplates using a Mosquito HV liquid handler (SPT Labtech). Final reaction conditions were as follows: enzyme concentrations of 17 µM and substrate concentrations of 0.1 mM were. Reaction microplates were sealed using a thermal heat sealer (Velocity11’s PlateLoc, Agilent Technologies) and incubated 24 hours at 25°C with agitation of 500 rpm.

Reactions were quenched with 4 µl of acetonitrile (AcN) 100% (v/v), followed by resealing the microplates, mixing of the plates for 5 minutes at 1000 rpm and centrifuging for 30 min at 6000 rpm. Microplates were stored at -20°C prior to analysis. High throughput screening for detection of substrate and/or product in each of the 480 reactions was performed by using Ultra Performance Liquid Chromatography (UPLC) using a Waters Acquity Ultra-Performance Liquid Chromatography (UPLC) system coupled with a SYNAPT G2 (HDMS) quadrupole time-of-flight (QToF) mass spectrometer (Waters Corp.). Liquid chromatography was performed by injecting 10 µl of each sample into an Acquity UPLC BEH C18 1.7 µm column at a flow rate of 0.5 mL/min. The mobile phase was prepared as a mixture of water with 0.1% formic acid (A) and ACN with formic acid 0.1% (B) with the following elution gradient: 0.00-6.00 min, 90-30% A; 6.00-6.70 min, 30-10% A; 6.70-7.00 min, 10-90% A; total run time was 7 minutes. Mass spectrometry was performed as follows: electrospray ionization data were obtained using the negative ion mode (ESI-) and a 2 kV capillary voltage; temperature of desolvation was 550 °C and source temperature was 120 °C; desolvation gas (nitrogen) flow was set to 850 L/Hr.

Data acquisition and processing were performed using MassLynx v4.2 (Waters Corp.). The MS profiles for each reaction were analysed to identify the degradation of substrate and/or the generation of product. Reactions where at least one of these two conditions could be observed were identified as positive hits in our high throughput screening and were classified according to the relative efficiency of the ancestral and modern glycosidases as described in Table S2.

### 4.4 | Enzymatic activity determinations

Hits where the ancestral glycosidase displayed a higher depletion of substrate and higher generation of expected flavonoid aglycon product than its modern counterpart were further characterised in upscaled enzymatic reactions. High molecular weight glycosylated flavonoids linarin, narirutin and hesperidin were identified as hits and used as substrates for the determination of reaction profiles and Michaelis-Menten curves for kinetic characterization. To quantify the concentration of product, we prepared standard curves for calibration by UPLC-MS using suitable dilutions with the de-glycosylated flavonoids expected to be released from each hit (acacetin, naringenin and hesperitin as products for linarin, narirutin and hesperidin, respectively). High purity flavonoids were diluted to prepare stock solutions of 0.1 M in DMSO and further diluted to prepare the standards for the curve.

First, a reaction profile was obtained by quantifying the amount of aglycon product generated by the ancestral and modern glycosidases at different times of reaction incubation with the substrates (see Figures S3-S5). Reactions were performed by incubating protein at 17 µM with 220 µM of each glycosylated flavonoid at 25 °C during different times and were quenched by adding 75 µl of the reaction solution to 225 µl of ACN 100% (v/v), followed by centrifugation for 15 mins at 14000 rpm 4 °C. Samples from the soluble fraction were extracted for further UPLC-MS analysis.

For initial rate calculations, reaction mixtures were incubated at 25°C for different times (10 mins to 3 hours) to determine a time at which that the amount of hydrolysed substrate was a small fraction of the total substrate and corresponded to the initial linear part of the reaction profiles. Measurements were corrected by a blank reaction performed under the same conditions. Reactions were quenched by adding 75 µl of the reaction solution to 225 µl of ACN 100% (v/v) and centrifuging for 15 mins at 14000 rpm 4 °C. Samples from the soluble fraction were extracted for further UPLC-MS analysis. For each of the glycoside flavonoids, initial rates were determined at several substrate concentrations within the range 0.05-3 mM. Catalytic parameters were determined by fitting the Michalis-Menten equation to the resulting profiles of initial rate *versus* substrate concentration.

In all cases, product formation determinations were performed with Ultra-High Performance Liquid Chromatography (UPLC) using a Waters Acquity H Class Ultra-Performance Liquid Chromatography (UPLC) system coupled with a Triwave Waters SYNAPT G2 and a high definition (HDMS) quadrupole time-of-flight (QToF) mass spectrometer (Waters Corp.). Liquid chromatography was performed by injecting 10 µl of each sample into an Acquity UPLC BEH ShieldRP18 1.7 µm column at a flow rate of 0.4 ml/min. The mobile phase was prepared as a mixture of water with 0.1% formic acid (A) and ACN with formic acid 0.1% (B) with an elution gradient as it follows: 0.00-9.00 min, 0-100% A; 9.00-10.00 min, stay 0% A; 10.00-12.00 min, 0-100% A; total run time 12 mins. Mass spectrometry was performed by using the following conditions: electrospray ionization data were obtained using the negative ion mode (ESI-) and a 2.6 kV capillary voltage. Temperature of desolvation was 500 °C and source temperature was 100 °C. Desolvation gas (nitrogen) flow was set to 1000 L/Hr. Data acquisition and processing were performed using MassLynx v4.2 (Waters Corp.).

### 4.5 | Molecular simulations

The multiple sequence alignment (MSA) required as input for the Boltz-1 calculations was generated using the GPU accelerated MMseqs2 software suite (Steinegger and Söding, 2017; Kallenborn et al., 2025). The enzyme sequences used are listed in the supplementary material as well as the SMILES strings for the substrates (Table S1). Default parameters were used for the Boltz-1 runs; 10 structures per run were produced for both the apo and the small-molecule complex structures.

MM-GBSA calculations available from the Schrödinger Prime protein structure prediction solution were used is used to estimate relative binding affinity for all the substrates of Figure 1 and all the enzymes considered in this study. Each of the 10 Boltz-1 predicted binding poses was used and we report always the average and standard deviations. Previous to each calculation the complexes were prepared with Schrödinger’s protein preparation module and the default parameters.

## Supporting Information

The authors give information on experimental procedures, as well as the sequences of the modern and the ancestral glycosidases.

## Acknowledgements

This research was supported by Grant PID2021-124534OB-100 (to J.M. S.-R.) funded by MICIU/AEI/10.13039/501100011033 and by FEDER/UE. K.Z. and P.S. were funded by the AstraZeneca postdoc programme. We are grateful to the technical staff of the High-Resolution Mass Spectrometry facilities at the “Centro de Instrumentación Científica - Universidad de Granada” for providing access to instrumentation and assistance with mass spectrometry experiments and data analysis.

## Data availability statement

The data that support the conclusions of this study are available from the authors upon reasonable request.

