## Supporting Information for "Ancestral *versus* modern substrate scope in family-1 glycosidases"

[3]Centro de Investigación en Tecnologías de la Información y las Telecomunicaciones (CITIC-UGR), Universidad de Granada, 18071-Granada, Spain.

[4]Departamento de Ciencias de la Computacion e Inteligencia Artificial, Escuela Tecnica Superior de Ingenierías Informática y de la Telecomunicacion, Universidad de Granada, 18071-Granada, Spain.

[5]Andalusian Research Institute in Data Science and Computational Intelligence (DaSCI), Universidad de Granada, 18071-Granada, Spain.

[6]Medicinal Chemistry, Research and Development, Respiratory and Immunology, BioPharmaceuticals R&D, AstraZeneca, 431 50 Gothenburg, Sweden.

[7]Early Chemical Development, Pharmaceutical Sciences, Biopharmaceuticals R&D, AstraZeneca, CB2 0AA Cambridge, United Kingdom.

**Correspondence**
Martin A. Hayes. Discovery Sciences, BioPharmaceuticals R&D AstraZeneca, 431 50 Gothenburg, Sweden..
Jose M. Sanchez-Ruiz. Departamento de Quimica Fisica. Facultad de Ciencias, Unidad de Excelencia de Quimica Aplicada a Biomedicina y Medioambiente (UEQ), Universidad de Granada, 18071 Granada, Spain..

**Table S1.** SMILE strings for the compounds of Figure 1

| Compound | CID | PubChem | LongName | SMILES |
|---|---|---|---|---|
| cmpnd1 | pNP-glu | 92930 | 4-nitrophenyl-b-D-glucopyranoside | C1=CC(=CC=C1[N+](=O)[O-])O[C@H]2[C@@H]([C@H]([C@@H]([C@H](O2)CO)O)O)O |
| cmpnd2 | pNP-gal | 65115 | 4-nitrophenyl-b-D-galactopyranoside | C1=CC(=CC=C1[N+](=O)[O-])O[C@H]2[C@@H]([C@H]([C@H]([C@H](O2)CO)O)O)O |
| cmpnd3 | MUB-gal | 93577 | 4-Methylumbelliferyl β-D-galactopyranoside | CC1=CC(=O)OC2=C1C=CC(=C2)O[C@H]3[C@@H]([C@H]([C@H]([C@H](O3)CO)O)O)O |
| cmpnd4 | RES-gal | 24899376 | Resorufin β-D-galactopyranoside | C/C(=C/C=C)/C(=O)N1CCOCC1 |
| cmpnd5 | FMG-gal | 128077 | Fluorescein mono-β-D-galactopyranoside | C1=CC=C2C(=C1)C(=O)OC23C4=C(C=C(C=C4)O)OC5=C3C=CC(=C5)O[C@H]6[C@@H]([C@H]([C@H]([C@H](O6)CO)O)O)O |
| cmpnd6 | RED-gal | 127428 | Chlorophenol Red-β-D-galactopyranoside | C1=CC=C2C(=C1)C(OS2(=O)=O)(C3=CC(=C(C=C3)O)Cl)C4=CC(=C(C=C4)O[C@H]5[C@@H]([C@H]([C@H]([C@H](O5)CO)O)O)O)Cl |
| cmpnd7 | LNR | 5317025 | Linarin | C[C@H]1[C@@H]([C@H]([C@H]([C@@H](O1)OC[C@@H]2[C@H]([C@@H]([C@H]([C@@H](O2)OC3=CC(=C4C(=C3)OC(=CC4=O)C5=CC=C(C=C5)OC)O)O)O)O)O)O)O |
| cmpnd8 | NRT | 442431 | Narirutin | C[C@H]1[C@@H]([C@H]([C@H]([C@@H](O1)OC[C@@H]2[C@H]([C@@H]([C@H]([C@@H](O2)OC3=CC(=C4C(=O)C[C@H](OC4=C3)C5=CC=C(C=C5)O)O)O)O)O)O)O)O |
| cmpnd9 | HPN | 10621 | Hesperidin | C[C@H]1[C@@H]([C@H]([C@H]([C@@H](O1)OC[C@@H]2[C@H]([C@@H]([C@H]([C@@H](O2)OC3=CC(=C4C(=O)C[C@H](OC4=C3)C5=CC(=C(C=C5)OC)O)O)O)O)O)O)O)O |

**Table S2**. Classification for each of the 291 substrates of the chemical library detected in LC-MS according to the activity profile of modern and ancestral glycosidase. No conversion was defined when <5% of substrate depletion and/or product generation was detected. Incomplete conversion was defined when 5-95% of substrate depletion and/or product generation was detected. Full conversion was defined when >95% of substrate depletion and/or product generation was detected.

| 291 substrates detected in LC-MS | ANCESTRAL | | | |
|---|---|---|---|---|
| MODERN | | NO CONVERSION | INCOMPLETE CONVERSION | FULL CONVERSION |
| | NO CONVERSION | **176** | **0** | **0** |
| | INCOMPLETE CONVERSION | **12** | **7** | **3** |
| | FULL CONVERSION | **12** | **26** | **55** |

**Table S3.** Molecular descriptors used to represent the relevant properties of the compounds in the screened library. The names of some descriptors include “scrn” meaning that the value corresponds to the whole molecule of the screened compound. Some descriptors include “agl” indicating that the value corresponds to the aglycon moiety exclusively. Strong correlations were found among certain sets of descriptors (see Figure S2). In these cases, only one member of the set was selected for decision-tree analysis. Selected descriptors are highlighted in bold.

| Descriptor Name | Molecular Description |
|---|---|
| **scrn_RingCount** | Numerical descriptor. Number of rings in the compound. |
| scrn_NumHeteroatoms | Numerical descriptor. Number of heteroatoms in the compound. |
| **scrn_NumHDonors** | Numerical descriptor. Number of hydrogen bond donors in the compound. |
| **scrn_NumHAcceptors** | Numerical descriptor. Number of hydrogen bond acceptors in the compound. |
| **scrn_NumAromaticHeterocycles** | Numerical descriptor. Number of aromatic heterocycles in the compound. |
| **scrn_NumAromaticCarbocycles** | Numerical descriptor. Number of aromatic carbocycles in the compound. |
| **scrn_NumAliphaticHeterocycles** | Numerical descriptor. Number of aliphatic heterocycles in the compound. |
| **scrn_NumAliphaticCarbocycles** | Numerical descriptor. Number of aliphatic carbocycles in the compound. |
| **scrn_Mol_Wt** | Numerical descriptor. Molecular weight of the compound in Dalton. |
| scrn_HeavyAtomCount | Numerical descriptor. Number of non-hydrogen atoms in the compound. |
| **scrn_Fraction_CSP3** | Numerical descriptor. Fraction of C atoms that are $sp^3$ hybridized in the compound. |
| **Monosaccharide** | Binary descriptor. Compound is a monosaccharide or not. |
| **Modified** | Binary descriptor. Classical sugar or deoxysugar, aminosugar, etc. |
| **M-1a** | Binary descriptor. Classical sugar with aliphatic radical or not. |
| **M-1b** | Binary descriptor. Classical sugar with aromatic radical or not. |
| **M-2a** | Binary descriptor. Hexose whose O is exchanged by a S with an aliphatic radical or not. |
| **M-2b** | Binary descriptor. Hexose whose O is exchanged by a S with an aromatic radical or not. |
| **M-3a** | Binary descriptor. Hexose whose O is exchanged by a N with an aliphatic.¡ radical or not |
| **M-3b** | Binary descriptor. Hexose whose O is exchanged by a N with an aromoatic radical or not |
| **M-4a** | Binary descriptor. Sugar is a pentose or not. |
| **M-4b** | Binary descriptor. Sugar is a pentose with the O is exchanged by a N or not. |
| **Anomeric_Stereo** | Categorical descriptor. Refers to the configuration of the anomeric carbon. It can be alpha, beta or unknown. |
| agl_RingCount | Numerical descriptor. Number of rings in the aglycon. |
| agl_NumHeteroatoms | Numerical descriptor. Number of heteroatoms in the aglycon. |
| agl_NumHDonors | Numerical descriptor. Number of hydrogen bond donors in the aglycon. |
| agl_NumHAcceptors | Numerical descriptor. Number of Hydrogen bond acceptors in the aglycon. |
| agl_NumAromaticHeterocycles | Numerical descriptor. Number of aromatic heterocycles in the aglycon. |
| agl_NumAromaticCarbocycles | Numerical descriptor. Number of aromatic carbocycles in the aglycon. |
| agl_NumAliphaticHeterocycles | Numerical descriptor. Number of aliphatic heterocycles in the aglycon. |
| agl_NumAliphaticCarbocycles | Numerical descriptor. Number of aliphatic carbocycles in the aglycon. |
| agl_Mol_Wt | Numerical descriptor. Molecular weight of the aglycon in Dalton. |
| agl_HeavyAtomCount | Numerical descriptor. Number of non-hydrogen atoms in the aglycon. |
| agl_Fraction_CSP3 | Numerical descriptor. Fraction of C atoms that are $sp^3$ hybridized in the aglycon. |

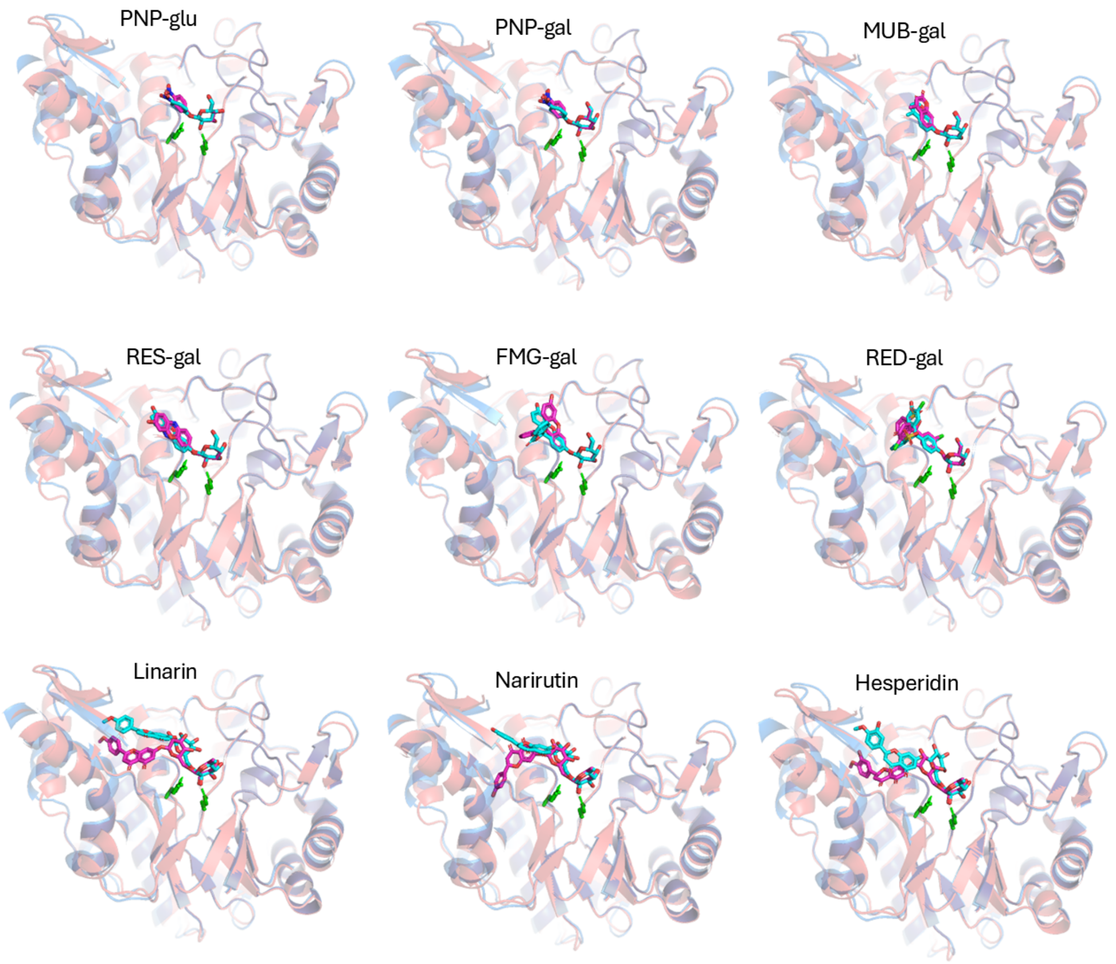


**Figure S1.** Protein structures with substrates docked, as predicted by Boltz-1. Modern is shown in blue and ancestral is shown in red. Figure 5 in the main text shows some representative examples, while additional illustrative examples are included here.

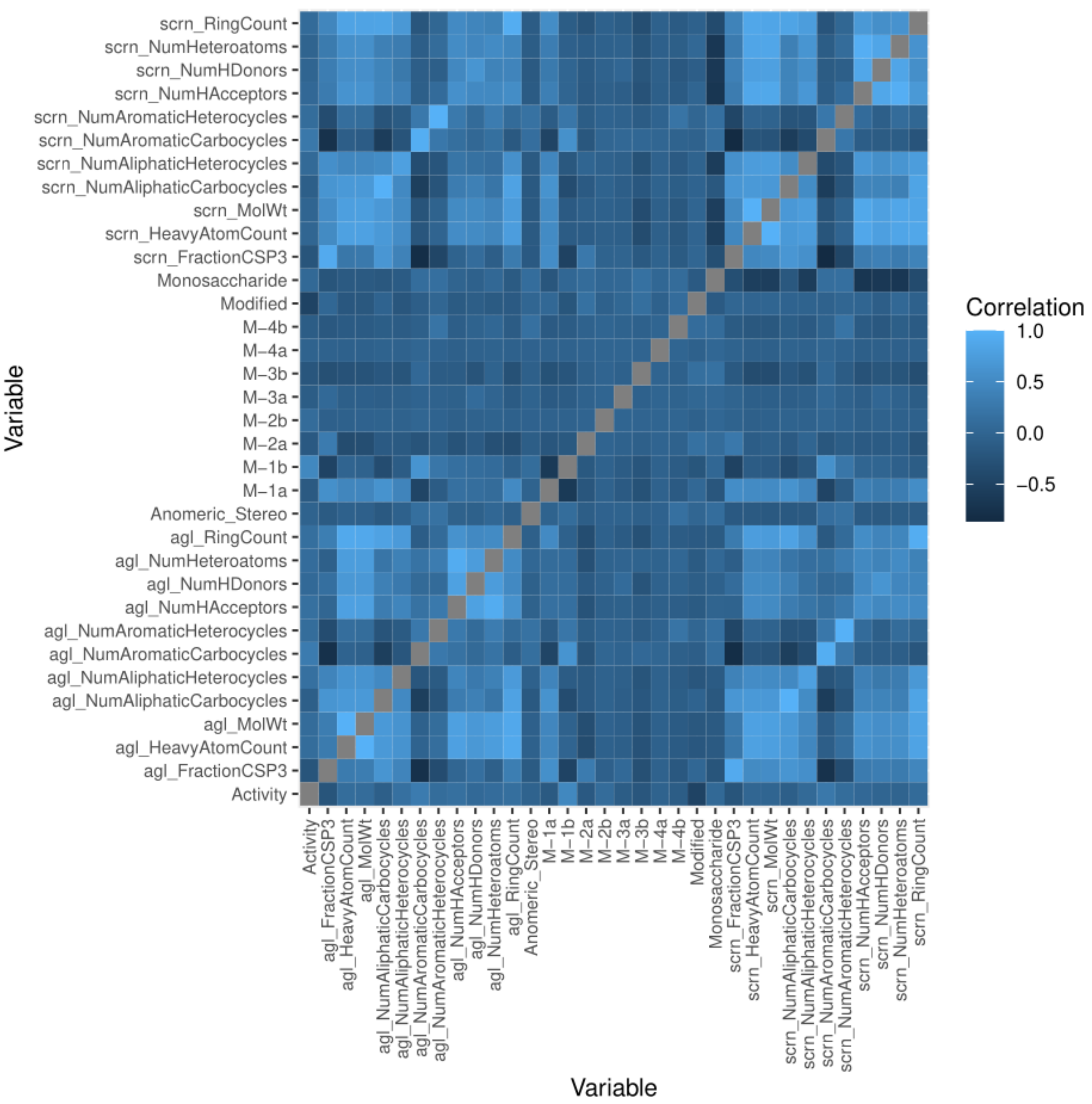


**Figure S2.** Pairwise correlation analysis of the molecular descriptors selected to represent the relevant properties of the compounds in the screened library. The colour code shown at the left describes the value of the Pearson correlation coefficient.

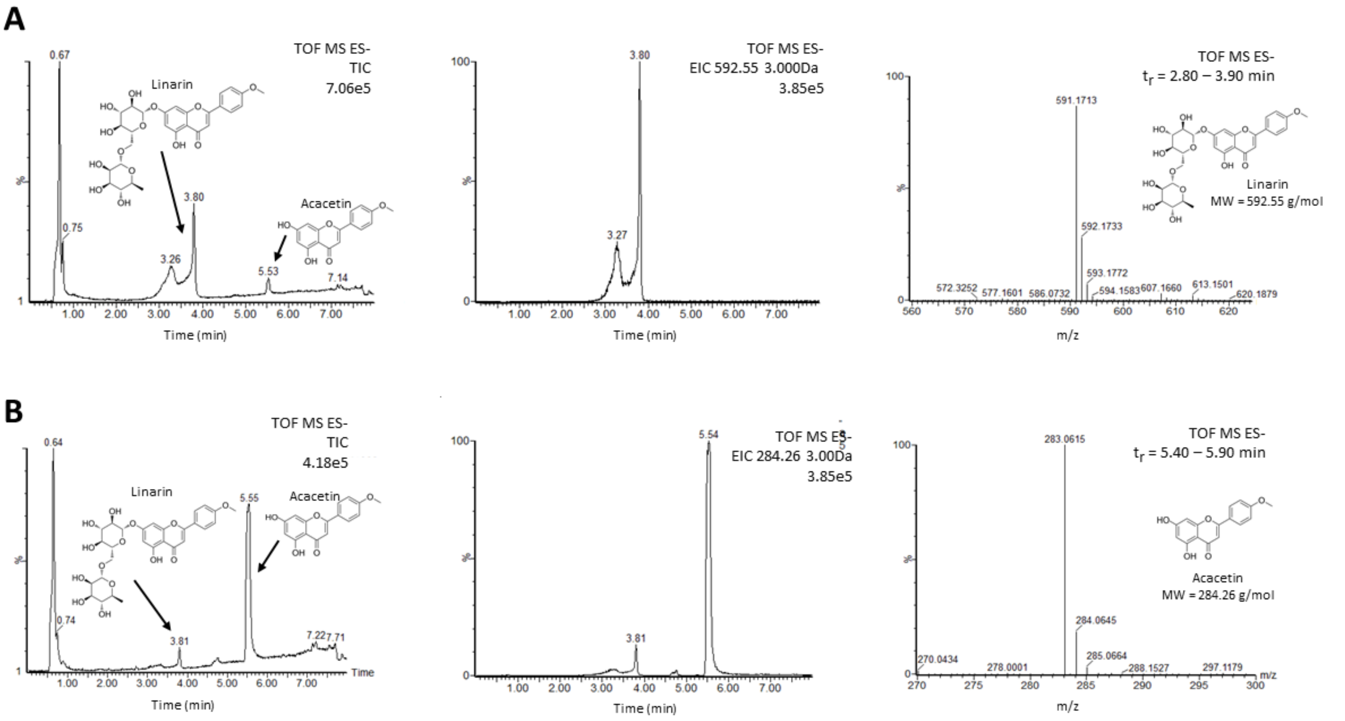


**Figure S3**. Mass spectrometry profiles for the hydrolysis of linarin. **A)** MS profiles for the hydrolysis of linarin after short incubation (2 min) with ancestral glycosidase. Left panel: Total ion count and peaks for both substrate (linarin) and deglycosylated flavonoid (acacetin) can be observed with retention times of 3.26 and 3.80 for linarin, and 5.53 for acacetin. Middle panel: Extracted ion count for the molecular weight of linarin (592.55 g/mol) confirms the presence of glycosylated flavonoid in the reaction. Right panel: UPLC-MS (QToF) data of the substrate peaks with $t_r$ = 2.80 – 3.90 min confirming the expected mass for linarin. **B)** MS profiles for the hydrolysis of linarin after long incubation (120 min) with ancestral glycosidase. Left panel: Total ion count and peaks for both substrate (linarin) and deglycosylated flavonoid (acacetin) can be observed with retention times of 3.81 for linarin, and 5.55 for acacetin. Almost full conversion of substrate can be observed. Middle panel: Extracted ion count for the molecular weight of acacetin (284.26 g/mol) confirms the presence of de-glycosylated flavonoid in the reaction. Right panel: UPLC-MS (QToF) data of the substrate peaks with $t_r$ = 5.40 – 5.90 min confirming the expected mass for acacetin.

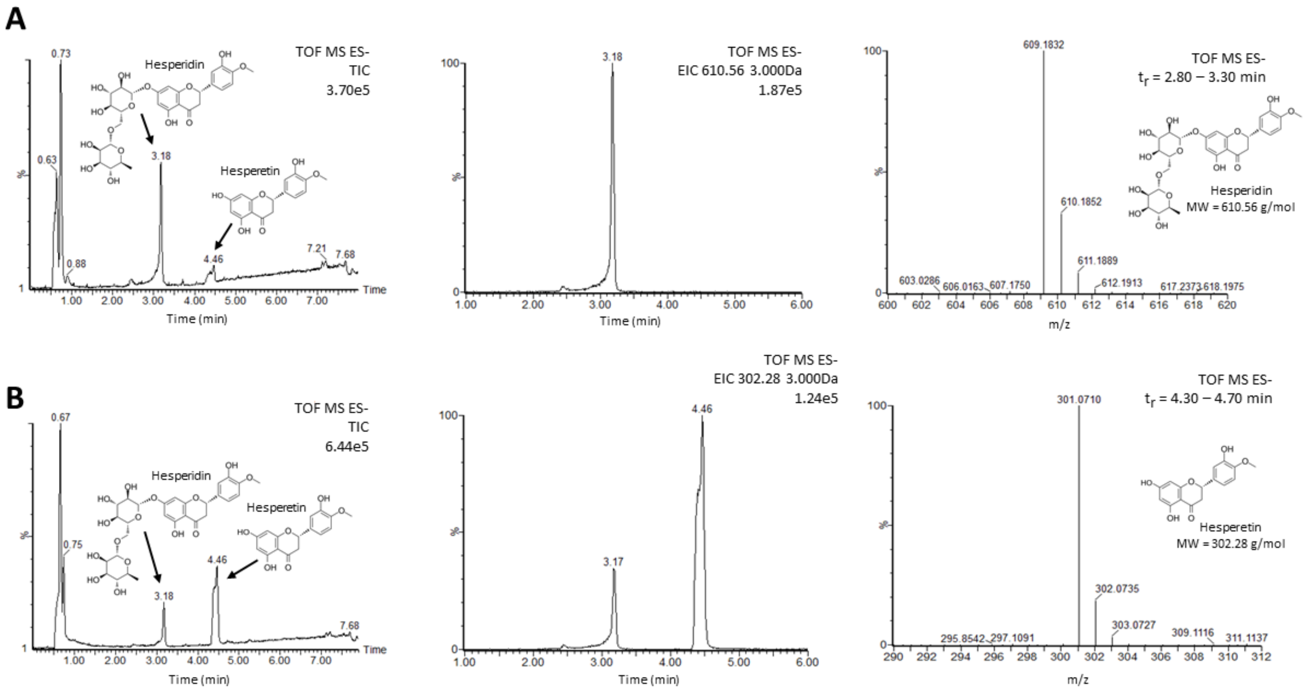


**Figure S4**. Mass spectrometry profiles for the hydrolysis of hesperidin. **A)** MS profiles for the hydrolysis of hesperidin after short incubation (4 min) with ancestral glycosidase. Left panel: Total ion count and peaks for both substrate (hesperidin) and deglycosylated flavonoid (hesperetin) can be observed with retention times of 3.18 for hesperidin, and 4.46 for hesperetin. Middle panel: Extracted ion count for the molecular weight of hesperidin (610.56 g/mol) confirms the presence of glycosylated flavonoid in the reaction. Right panel: UPLC-MS (QToF) data of the substrate peaks with $t_r$ = 2.80 – 3.30 min confirming the expected mass for hesperidin. **B)** MS profiles for the hydrolysis of hesperidin after long incubation (240 min) with ancestral glycosidase. Left panel: Total ion count and peaks for both substrate (hesperidin) and deglycosylated flavonoid (hesperetin) can be observed with retention times of 3.81 for hesperidin, and 4.46 for hesperetin. Almost full conversion of substrate can be observed. Middle panel: Extracted ion count for the molecular weight of hesperetin (302.28 g/mol) confirms the presence of de-glycosylated flavonoid in the reaction. Right panel: UPLC-MS (QToF) data of the substrate peaks with $t_r$ = 4.30 – 4.70 min confirming the expected mass for hesperetin.

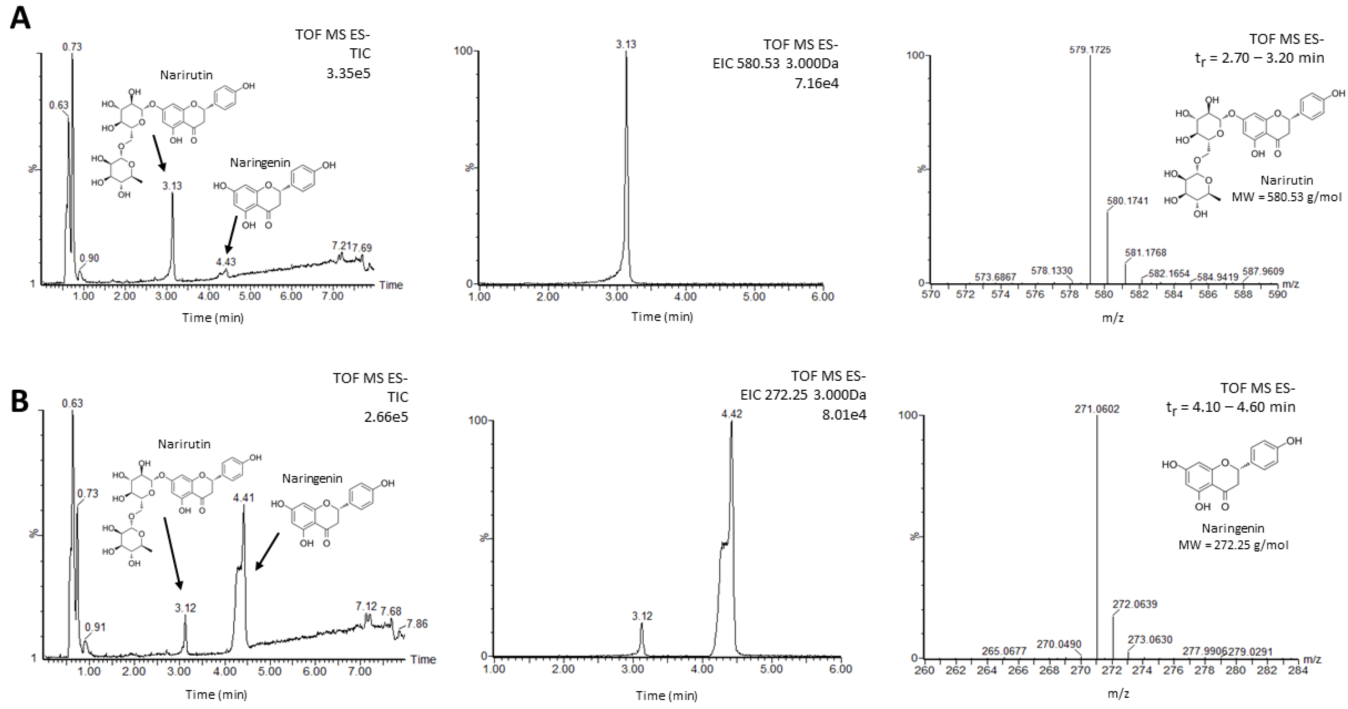


**Figure S5**. Mass spectrometry profiles for the hydrolysis of narirutin. **A)** MS profiles for the hydrolysis of narirutin after short incubation (8 min) with ancestral glycosidase. Left panel: Total ion count and peaks for both substrate (narirutin) and deglycosylated flavonoid (naringenin) can be observed with retention times of 3.13 for narirutin, and 4.43 for naringenin. Middle panel: Extracted ion count for the molecular weight of naringenin (580.53 g/mol) confirms the presence of glycosylated flavonoid in the reaction. Right panel: UPLC-MS (QToF) data of the substrate peaks with $t_r$ = 2.70 – 3.20 min confirming the expected mass for narirutin. **B)** MS profiles for the hydrolysis of narirutin after long incubation (240 min) with ancestral glycosidase. Left panel: Total ion count and peaks for both substrate (narirutin) and deglycosylated flavonoid (naringenin) can be observed with retention times of 3.12 for narirutin, and 4.41 for naringenin. Almost full conversion of substrate can be observed. Middle panel: Extracted ion count for the molecular weight of naringenin (272.25 g/mol) confirms the presence of de-glycosylated flavonoid in the reaction. Right panel: UPLC-MS (QToF) data of the substrate peaks with $t_r$ = 4.10 – 4.60 min confirming the expected mass for naringenin.

### SEQUENCES OF MODERN AND ANCESTRAL GLYCOSIDASE

#### Sequences

The sequence of the ancestrally reconstructed enzyme is available from the RCSB PDB entry 6Z1H, chain A. The sequence of the modern GH1 is available from the UNIPROT entry B8CYA8.

```
>6Z1H_A

MTQTAAKSLKFPKDFLWGAATAAYQIEGAANEDGRGPSIWDTFSHTPGKVHNGDNGDVACDHYHRYKEDV
ELMKELGLNAYRFSISWPRILPEGEGKVNQKGLDFYNNLIDELLENGIEPFVTLYHWDLPQALQDKGGWE
NRETVDAFAEYARVCFERFGDRVKYWITFNEPNVFAVLGYLSGVHPPGMKDLKKAFRAAHNLLLAHARAV
KAYREISQNGQIGITLNLSPVYPASDNEEEDKAAAERADQFNNWFLDPIFKGKYEHMLERLGEQIAANGG
ELPEITDEMEILSASLDFIGLNYYTSNLVRANPNSGSSSVKPPDLPRTDMGWEIYPEGLYDLLKRIHEKY
NLPIYITENGMAVDDEVEDGAVHDTNRIDYLKEHLEAVHKAIEEGVNVRGYFVWSLMDNFEWANGYSKRF
GLIYVDYKTQKRTPKKSAYWYREVIKSNGLELEHHHHHH

>B8CYA8

MAKIIFPEDFIWGAATSSYQIEGAFNEDGKGESIWDRFSHTPGKIENGDTGDIACDHYHLYREDIELMKE
IGIRSYRFSTSWPRILPEGKGRVNQKGLDFYKRLVDNLLKANIRPMITLYHWDLPQALQDKGGWTNRDTA
KYFAEYARLMFEEFNGLVDLWVTHNEPWVVAFEGHAFGNHAPGTKDFKTALQVAHHLLLSHGMAVDIFRE
EDLPGEIGITLNLTPAYPAGDSEKDVKAASLLDDYINAWFLSPVFKGSYPEELHHIYEQNLGAFTTQPGD
MDIISRDIDFLGINYYSRMVVRHKPGDNLFNAEVVKMEDRPSTEMGWEIYPQGLYDILVRVNKEYTDKPL
YITENGAAFDDKLTEEGKIHDEKRINYLGDHFKQAYKALKDGVPLRGYYVWSLMDNFEWAYGYSKRFGLI
YVDYENGNRRFLKDSALWYREVIEKGQVEAN
```

#### Alignment

The sequence alignment below was performed with clustalW 2.1, with standard parameters. The C-terminal poly-histidine tag in the ancestrally reconstructed GH1 was omitted.

```
6Z1H_A  MTQTAAKSLKFPKDFLWGAATAAYQIEGAANEDGRGPSIWDTFSHTPGKVHNGDNGDVAC
B8CYA8  -----MAKIIFPEDFIWGAATSSYQIEGAFNEDGKGESIWDRFSHTPGKIENGDTGDIAC
               .: **:**:*****::****** ****:* **** *******:.***.**:**

6Z1H_A  DHYHRYKEDVELMKELGLNAYRFSISWPRILPEGEGKVNQKGLDFYNNLIDELLENGIEP
B8CYA8  DHYHLYREDIELMKEIGIRSYRFSTSWPRILPEGKGRVNQKGLDFYKRLVDNLLKANIRP
        **** *:**:*****:*:.:**** *********:*:*********:.*:*:**: .*.*

6Z1H_A  FVTLYHWDLPQALQDKGGWENRETVDAFAEYARVCFERFGDRVKYWITFNEPNVFAVLGY
B8CYA8  MITLYHWDLPQALQDKGGWTNRDTAKYFAEYARLMFEEFNGLVDLWVTHNEPWVVAFEGH
        ::***************** **:*.. ******: **.*.. *. *:*.*** *.*. *:
```

```
6Z1H_A  LSGVHPPGMKDLKKAFRAAHNLLLAHARAVKAYREISQNGQIGITLNLSPVYPASDNEEE

B8CYA8  AFGNHAPGTKDFKTALQVAHHLLLSHGMAVDIFREEDLPGEIGITLNLTPAYPAGDSEKD

        * *.** **:*.*::.**:***:*. **. :** .  *:*******:*.***.*.*::


6Z1H_A  DKAAAERADQFNNWFLDPIFKGKYEHMLERLGEQIAANGGELPEITDEMEILSASLDFIG

B8CYA8  VKAASLLDDYINAWFLSPVFKGSYPEELHHIYEQ---NLGAFTTQPGDMDIISRDIDFLG

        ***:   * :* ***.*:***.* . *.:: **   * * :.  ..:*:*:* .:**:*


6Z1H_A  LNYYTSNLVRANPNS---GSSSVKPPDLPRTDMGWEIYPEGLYDLLKRIHEKYN-LPIYI

B8CYA8  INYYSRMVVRHKPGDNLFNAEVVKMEDRPSTEMGWEIYPQGLYDILVRVNKEYTDKPLYI

        :***:  :** :*..   .:. **  * * *:*******:****:* *::::*.  *:**


6Z1H_A  TENGMAVDDEV-EDGAVHDTNRIDYLKEHLEAVHKAIEEGVNVRGYFVWSLMDNFEWANG

B8CYA8  TENGAAFDDKLTEEGKIHDEKRINYLGDHFKQAYKALKDGVPLRGYYVWSLMDNFEWAYG

        **** *.**:: *:* :** :**:** :*:: .:**:::** :***:*********** *


6Z1H_A  YSKRFGLIYVDYKT-QKRTPKKSAYWYREVIKSNGLELE

B8CYA8  YSKRFGLIYVDYENGNRRFLKDSALWYREVIEKGQVEAN

        ************:. ::*  *.** ******:.. :* :
```

**SEQUENCES OF ENZYME VARIANTS**

>gh1-a2m

MTQTAAKSLKFPKDFLWGAATAAYQIEGAANEDGRGPSIWDTFSHTPGKVHNGDN
GDVACDHYHRYKEDVELMKELGLNAYRFSISWPRILPEGEGKVNQKGLDFYNNLI
DELLENGIEPFVTLYHWDLPQALQDKGGWENRETVDAFAEYARVCFERFGDRVKY
WITFNEPWVFAFEGYASGVHPPGMKDFKKAFRAAHNLLLAHARAVKAYREISQNG
QIGITLNLTPVYPASDNEEEDKAAAERADQINNWFLDPIFKGKYEHMLERLGEQI
AANGGELPEITDEMEILSASLDFIGLNYYSSMLVRANPNSGSSSVKPPDLPRTDM
GWEIYPEGLYDLLKRIHEKYNLPIYITENGMAVDDEVEDGAVHDTNRIDYLKEHL
EAVHKAIEEGVNVRGYFVWSLMDNFEWANGYSKRFGLIYVDYKTQKRTPKKSAYW
YREVIKSNGLELE

>gh1-a2m2

MTQTAAKSLKFPKDFLWGAATAAYQIEGAANEDGRGPSIWDTFSHTPGKVHNGDN
GDVACDHYHRYKEDVELMKELGLNAYRFSISWPRILPEGEGKVNQKGLDFYNNLI
DELLENGIEPFVTLYHWDLPQALQDKGGWENRETVDAFAEYARVCFERFGDRVKY
WITFNEPWVFAFEGYAFGVHPPGMKDFKKAFRAAHNLLLAHARAVKAYREISQNG
QIGITLNLTPVYPASDNEEEDKAAAELADQINNWFLDPIFKGKYEHELERLYEQI
AANGGELPEITDEMEILSASLDFIGLNYYSSMLVRANPNSGSSSVKPPDLPRTDM
GWEIYPEGLYDLLKRIHEKYNLPIYITENGMAVDDEVEDGAVHDTNRIDYLKEHL
EAVHKAIEEGVNVRGYFVWSLMDNFEWANGYSKRFGLIYVDYKTQKRTPKKSAYW
YREVIKSNGLELE

>gh1-m2a

MAKIIFPEDFIWGAATSSYQIEGAFNEDGKGESIWDRFSHTPGKIENGDTGDIAC
DHYHLYREDIELMKEIGIRSYRFSTSWPRILPEGKGRVNQKGLDFYKRLVDNLLK
ANIRPMITLYHWDLPQALQDKGGWTNRDTAKYFAEYARLMFEEFNGLVDLWVTHN
EPNVVAVLGHLFGNHAPGTKDLKTALQVAHHLLLSHGMAVDIFREEDLPGEIGIT
LNLSPAYPAGDSEKDVKAASLLDDYFNAWFLSPVFKGSYPEELHHIYEQNLGAFT
TQPGDMDIISRDIDFLGINYYTRNVVRHKPGDNLFNAEVVKMEDRPSTEMGWEIY
PQGLYDILVRVNKEYTDKPLYITENGAAFDDKLTEEGKIHDEKRINYLGDHFKQA
YKALKDGVPLRGYYVWSLMDNFEWAYGYSKRFGLIYVDYENGNRRFLKDSALWYR
EVIEKGQVEAN

>gh1-m2a2

MAKIIFPEDFIWGAATSSYQIEGAFNEDGKGESIWDRFSHTPGKIENGDTGDIAC
DHYHLYREDIELMKEIGIRSYRFSTSWPRILPEGKGRVNQKGLDFYKRLVDNLLK

ANIRPMITLYHWDLPQALQDKGGWTNRDTAKYFAEYARLMFEEFNGLVDLWVTHN
EPNVVAVLGHLSGNHAPGTKDLKTALQVAHHLLLSHGMAVDIFREEDLPGEIGIT
LNLSPAYPAGDSEKDVKAASLRDDYFNAWFLSPVFKGSYPEMLHHIGEQNLGAFT
TQPGDMDIISRDIDFLGINYYTRNVVRHKPGDNLFNAEVVKMEDRPSTEMGWEIY
PQGLYDILVRVNKEYTDKPLYITENGAAFDDKLTEEGKIHDEKRINYLGDHFKQA
YKALKDGVPLRGYYVWSLMDNFEWAYGYSKRFGLIYVDYENGNRRFLKDSALWYR
EVIEKGQVEAN
